# Context sensitivity across multiple time scales with a flexible frequency bandwidth

**DOI:** 10.1101/2020.06.08.141044

**Authors:** Tamar I. Regev, Geffen Markusfeld, Leon Y. Deouell, Israel Nelken

## Abstract

Everyday auditory streams are complex, including spectro-temporal content that varies at multiple time scales. Using EEG, we investigate the sensitivity of human auditory cortex to the content of past stimulation in unattended sequences of equiprobable tones. In 3 experiments including 82 participants overall, we found that neural responses measured at different latencies after stimulus onset were sensitive to frequency intervals computed over distinct time scales. Importantly, early responses were sensitive to a longer history of stimulation than later responses. To account for these results, we tested a model consisting of neural populations with frequency-specific but broad tuning that undergo adaptation with exponential recovery. We found that the coexistence of neural populations with distinct recovery rates can explain our results. Furthermore, the adaptation bandwidth of these populations depended on spectral context – it was wider when the stimulation sequence had a wider frequency range. Our results provide electrophysiological evidence as well as a possible mechanistic explanation for dynamic and multi-scale context-dependent auditory processing in the human cortex.

**SIGNIFICANCE STATEMENT:** It has become clear that brain processing of sensory stimuli depends on their temporal context, but context can be construed at time scales from the recent millisecond to life-long. How do different contextual time scales affect sensory processing? We show that auditory context is integrated across at least two separate time scales, and that at both of these time scales responses dynamically adapt to a varying frequency stimulation range. Using computational modeling, we develop a rigorous methodology to estimate the time and frequency scales of context integration for separate response components. Our robust results replicated across 3 EEG experiments, and contribute to the understanding of neural mechanisms supporting complex and dynamic context integration.

## INTRODUCTION

To function efficiently, sensory systems should interpret incoming stimuli in the context in which they are embedded. Contextual effects in audition are often studied using sound sequences that have some regularity, which is infrequently violated by ‘deviant’ sounds. These studies show that in humans, auditory event-related potentials depend on the preceding statistics of the sequence (Sussman, 2007; Garrido et al., 2013; Herrmann et al., 2015). However, processing of context is important for any stimulus sequence structure and not just for detecting regularities or change thereof. Here, we generalize contextual effects beyond deviance processing, using human EEG recordings, and modelling.

One way of efficiently representing context is by summary statistics of past stimulation. Indeed, humans can reliably report the mean pitch of several pure tones (Albrecht et al., 2012; Piazza et al., 2013), and sound textures are represented using time-averaged statistics (McDermott et al., 2013). As the environment constantly changes, estimating summary descriptors dynamically may optimize information transmission (Brenner et al., 2000; Fairhall et al., 2001). Across species and modalities, neuronal input-output functions scale with statistical properties of the stimulus distribution including mean (Dunn and Rieke, 2006; Nagel and Doupe, 2006; Dean et al., 2008; Dahmen et al., 2010), variance (Blake and Merzenich, 2002; Maravall et al., 2007; Rabinowitz et al., 2011; Herrmann et al., 2013, 2014, 2015) or higher order moments (Kvale and Schreiner, 2004; Herrmann et al., 2018, 2020). The time scales over which context influences neural activity vary widely, from tens of milliseconds to minutes (Fairhall et al., 2001; Khouri and Nelken, 2015). Presumably, this variation is necessary because the natural auditory environment contains relevant information at all of these time scales. Sensitivity of auditory neural responses to regularities established across multiple time scales was reported in single A1 neurons in animals (Ulanovsky et al., 2004), human MEG (Maheu et al., 2019) or the EEG components MMN and P2 (Costa-Faidella et al., 2011). However, all of the above studies concentrated on deviance detection.

We investigated context-dependent auditory processing not involving deviance-detection mechanisms by measuring EEG responses to tone sequences in which all stimuli were equi-probable and task-irrelevant. First, we address data from the control conditions of two experiments (1 and 2) previously published (Regev et al., 2019). We observed in these data that the N1 and P2 event-related EEG components (peaking ~100 and ~180ms following stimulus presentation, respectively) were sensitive to past frequencies across distinct time scales. Previous studies established that the time scale of N1 sensitivity is longer than ~1s (Zacharias et al., 2012; Okamoto and Kakigi, 2014; Herrmann et al., 2016). Further, the context sensitivity of the N1 component was well explained by adaptation models, but the P2 component was not (Herrmann et al., 2013). Importantly, the latter study used a predetermined adaptation time constant to model both N1 and P2. Here, we use a frequency-specific adaptation model (Herrmann et al., 2013, 2014, 2015) to explain our previous results as well as a new dataset (Experiment 3), but instead of using a predetermined time constant, we develop a rigorous methodology to quantitatively estimate the effective time scales of each component from the data. Consequently, we show that auditory responses at different latencies (i.e., the latencies of the N1 and P2 components) can be explained by the same mechanism of frequency-specific adaptation, but with different adaptation time scales. Notably, the earlier component (N1) was sensitive to a longer history of stimulation than the later component (P2). Furthermore, previous studies suggested that the adaptation bandwidth of N1, but not of P2, dynamically adapts to the range of frequencies presented in the stimulus stream (Herrmann et al., 2013). Therefore, in Experiment 3 we manipulated the overall range of frequencies in the sequences. We replicated the dynamic spectral-context-dependent adaptation bandwidth of N1, and generalized it to the P2 latency. We conclude that this type of adaptation, with varying bandwidths and time scales, is a general mechanism shaping auditory event-related potentials.

## METHODS

### Participants

Eighty-nine healthy adults participated in all 3 experiments - 25 musicians, 29 musicians, and 35 non-musicians in Experiments 1, 2 and 3, respectively. The reason musicians participated in Experiments 1 and 2 is not relevant to the current study and is explained in (Regev et al., 2019). Participants were recruited from The Hebrew University of Jerusalem, from Bezalel Academy of Arts and Design or the Jerusalem Academy of Music and Dance, and could either receive 40 NIS (~12 US$) per hour or course credit for participation in the experiment. The data of 7 participants (4, 1 and 2 from Experiments 1, 2 and 3, respectively) were excluded due to technical difficulties with the recording or excessive rates of artifacts. The analysis therefore included the data of 82 participants – 21 in Experiment 1 (7 female, mean age = 29.2 years, SD = 9 years), 28 in Experiment 2 (15 female, mean age = 24.6 years, SD = 3.6 years), and 33 participants in Experiment 3 (19 females, mean age = 24.6, SD = 2.9 years old). Three more participants (1 from Experiment 2 and 2 from Experiment 3) were later excluded from data analysis due to unclear auditory responses, resulting in 79 participants overall in the critical analysis as explained further in the methods section *Data Processing*. All participants self-reported normal hearing and no history of neurological disorders. The experiment was approved by the ethical committee of the faculty of social science at The Hebrew University of Jerusalem, and informed consent was obtained after the experimental procedures were explained.

### Stimuli and apparatus

In all experiments, participants were seated in a dimly lit, sound-attenuated, and echo-reduced chamber (C-26, Eckel) in front of a 17-inch CRT monitor (100-Hz refresh rate), at a viewing distance of about 90 cm. The screen was concealed by a black cover, with a rectangular window in the middle (14 by 8.5 cm), through which they viewed the visual display. Auditory stimuli were presented through headphones (Sennheiser HD25, having a relatively flat frequency response function in the range of frequencies used in the experiment) that was placed over the EEG cap. The experiment was run using the Psychophysics toolbox (Brainard, 1997) for MATLAB (version 2013b, MathWorks) running on a 32-bit Windows XP system. Auditory stimuli were synthesized using MATLAB. The experiment included only pure tones, each of 100 milliseconds (ms) duration with a 30 ms linear rise and fall ramps. Stimuli were presented at a sound pressure level that was comfortable for the participants. At the beginning of the experiment each participant adjusted the relative amplitudes of each individual tone, such that all tones had approximately the same subjective loudness.

### Experiment design

Participants viewed silent black and white films while tones were presented to them through headphones. The participants could choose either “The Artist” (Michel Hazanavicius, 2011) or “The Kid” (Charlie Chaplin, 1921), both silent movies. The participants were instructed to ignore the sounds. Each tone sequence in all 3 experiments was comprised of pure tones of 5 different frequencies. The specific frequencies varied between block types (see Figure 1, upper row, for an illustration of the stimuli, and a detailed description below). To create the sequences, random permutations of the 5 tones were concatenated successively. If the first tone of the next random permutation was the same as the last tone of the previous permutation, the order of tones in next permutation was reversed. As a result, the order of the tones was random with three constraints: i) each tone occurred exactly in 20% of the sound presentations, ii) a repetition of the same frequency never occurred, and iii) two successive presentations of the same tone frequency were separated by no more than 8 other sounds, imposing a substantial uniformity of tone occurrences over time.

**Figure 1.**
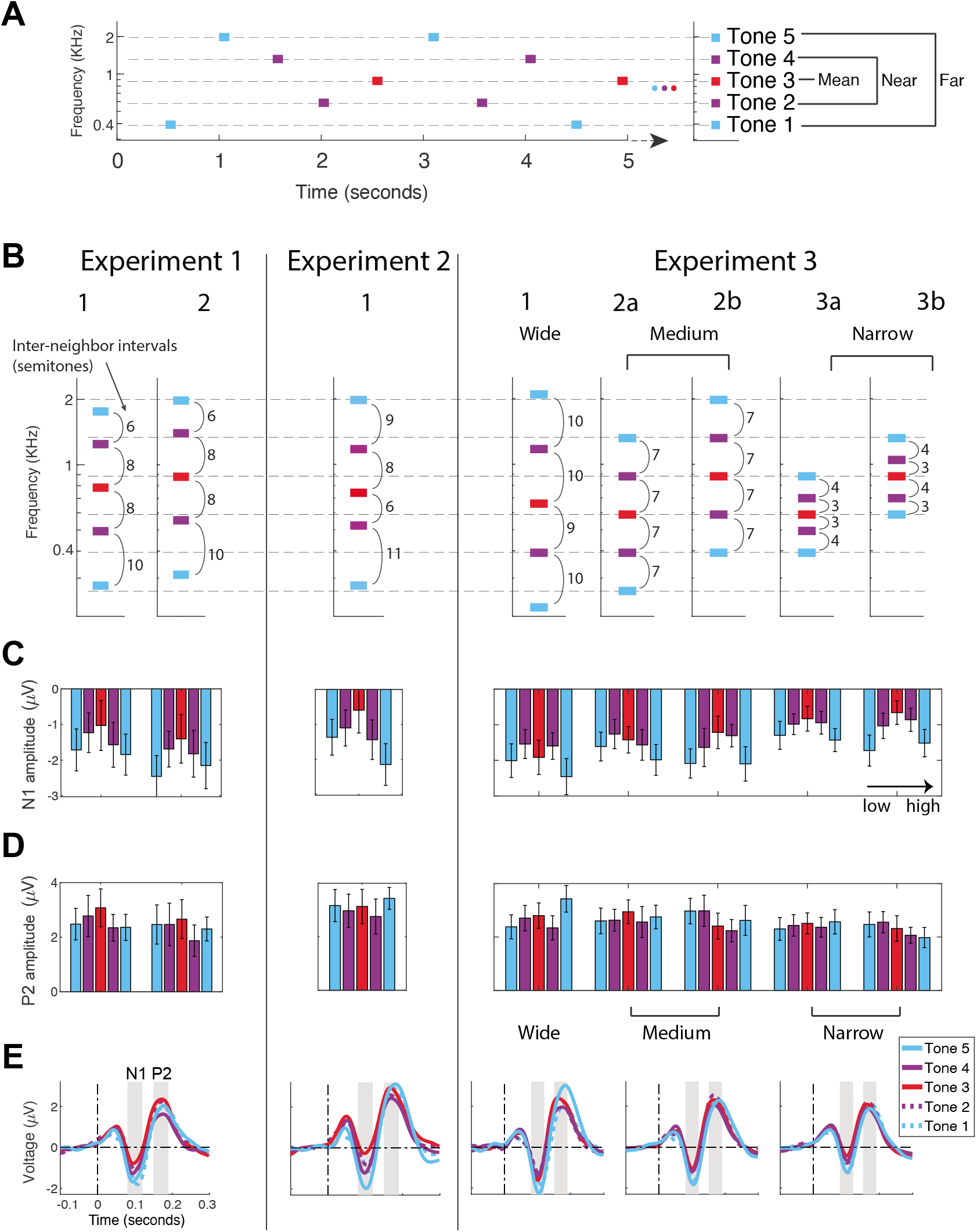
N1 but not P2 is sensitive to long-term context. **A** – An example segment of a tone sequence in the experiment (from block type 2b of Experiment 3). **B** – Stimuli used in all experiments and conditions. Intervals between neighboring tones on the frequency axis (inter-neighbor intervals) are displayed in semitones. **C** – Mean and 95% confidence intervals (across participants) of N1 peak amplitudes. For each block type the 5 bars correspond to tones 1, 2, 3, 4 and 5 (lowest to highest) from left to right and the bar colors match the color scheme in panels A and B. **D** – Same as C for peak amplitude of P2. **E** – Event-related potentials (ERPs) for tones 1 to 5 (low to high frequency), calculated for each experiment. For Experiment 3, ERPs are plotted for each frequency range, pooling together block types 2a+2b and 3a+3b.

#### Experiment 1

Two block types from Experiment 1 of Regev et al. (2019) (which served as control conditions in the previous study) were used for the current study. Condition 1 included 5 tones: Db4, B4, G5, Eb6 and A6 (277.2, 493.9, 784, 1244.5 and 1760 Hz, respectively). Hence, the tones spanned 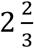 octaves (32 semitones) and the inter-neighbor intervals (frequency intervals between adjacent tones on the sequence-specific frequency axis, see Figure 1B) were 10, 8, 8 and 6 semitones from low to high frequency. The mean frequency (computed on the logarithmic frequency axis) was that of G5 – 784 Hz. Condition 2 included: Eb4, Db5, A5, F6 and B6 (311.1, 554.4, 880, 1397 and 1975.5 Hz, respectively). These were similar to the tones in condition 1, but all shifted 2 semitones down. Three blocks of each condition were presented and their order was counterbalanced between participants. Each block included 500 trials, 100 of each specific tone. This resulted in 300 trials for each specific tone in each condition. The tones were presented with an SOA (stimulus onset asynchrony, i.e., the time interval between the onsets of two consecutive stimuli) of either 450 or 550 ms, randomly (average SOA was 500 ms). As a result, each block lasted 250 s, and there were at least 30 s of silence between the blocks (at the participant discretion). See Figure 1B for illustration of stimuli.

#### Experiment 2

One block type from Experiment 2 of Regev et al. (2019) (which served as a control condition in the previous study) was used for the current study. This condition included 5 tones with equal probabilities: Db4, C5, F#5, D6 and B6 (277.9, 523.2, 740, 1174.7, and 1975.6 Hz, respectively). Hence, the tones spanned 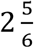 octaves (34 semitones) and the inter-neighbor intervals were 11, 6, 8 and 9 semitones from low to high frequency. The mean frequency (computed on the logarithmic frequency axis) was that of F#5 – 740 Hz. Two blocks of this condition were presented. Each block included 550 trials presented with an SOA of 400 ms. This resulted in 220 trials for each specific tone. Each block lasted 220 s, and there were at least 30 s of silence between the blocks (at the participant discretion). See Figure 1B for illustration of stimuli.

#### Experiment 3

In this experiment we manipulated the overall range of frequencies in the sequences (Table 1). Five block types were used. Block 1 with a wide range of frequencies (Wide; 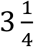 octaves between the lowest and highest tone, 39 semitones), Blocks 2a and 2b with a medium range (Medium; 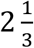 octaves, 28 semitones), and blocks 3a and 3b with a narrow range (Narrow; 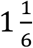 octaves, 14 semitones). The inter-neighbor intervals were 9.75, 7 and 3.5 semitones in the Wide, Medium and Narrow range conditions. For the Medium and Narrow conditions, we designed 2 different block types (a and b), transposed by 7 semitones, in order to generalize the results beyond the specific mean or range of frequencies (Figure 1B). Each block type was presented 3 times so the experiment consisted of 15 blocks. The order of the blocks was randomized for each participant separately, with the constraint that successive blocks had to be of different types. In every block 540 tones were presented in total, resulting in 324 trials overall for each tone frequency in each block type.

**Table 1.** Description of stimuli in Experiment 3. Tone properties are listed from low to high.

| Condition | 1 (wide) | 2a (medium) | 2b (medium) | 3a (narrow) | 3b (narrow) |
| --- | --- | --- | --- | --- | --- |
| Notes | A3, G4, E5, D6, C7 | C4, G4, D5, A5, E6 | G4, D5, A5, E6, B6 | G4, B4, D5, F5, A5 | D5, F5, A5, C6, E6 |
| Frequencies (Hz) | 220, 392, 659.26, 1174.7, 2093 | 261.6, 392, 587.33, 880, 1318 | 392, 587.3, 880, 1318, 1975.5 | 392, 493.88, 587.33, 698.46, 880 | 493.88, 698.46, 880, 1046.5, 1318 |
| Inter-neighbor intervals* (low to high, semitones) | 10, 9, 10, 10 | 7,7,7,7 | 7,7,7,7 | 4,3,3,4 | 3,4,3,4 |
| Mean frequency (note, Hz) | F5, 698.46 | D5, 587.33 | A5, 880 | D5, 587.33 | A5, 880 |
| Range (octaves) | $3\frac{1}{4}$ | $2\frac{1}{3}$ | $2\frac{1}{3}$ | $1\frac{1}{6}$ | $1\frac{1}{6}$ |
| Range (semitones) | 39 | 28 | 28 | 14 | 14 |
| Mean Inter-neighbor interval (semitones) | 9.75 | 7 | 7 | 3.5 | 3.5 |

\*Mean interval between nearby tones on the sequence-specific frequency scale, Figure 1B

The frequencies of all tones were taken from the C major scale, in order to prevent a situation in which one of the tones would become harmonically deviant and therefore result in stronger ERP responses (Poulin-Charronnat et al., 2006; Koelsch, 2009).

The SOA was randomly set to one of 5 possible values: 450, 475, 500, 525 or 550 ms. As a result, the duration of each block was about 270 s (4.5 minutes), and there were at least 45 seconds of silence between blocks (at the participant discretion). In total, the EEG was recorded for approximately an hour and a half.

### EEG recording and preprocessing

EEG was recorded from 64 pre-amplified Ag/AgCl electrodes using an Active 2 system (BioSemi, the Netherlands), mounted on an elastic cap according to the extended 10-20 system (https://www.biosemi.com/pics/cap_64_layout_medium.jpg), with the addition of two mastoid electrodes and a nose electrode. Horizontal electrooculogram (EOG) was recorded from electrodes placed at the outer canthi of the right and left eyes. Vertical EOG was recorded from electrodes placed below the center of both eyes and above the center of the right eye. The EEG signal was sampled at a rate of 512 Hz (24 bits/channel) with an online antialiasing low-pass filter set at one fifth of the sampling rate, and stored for offline analysis.

EEG preprocessing was conducted using BrainVision Analyzer 2.0 (Brain Products), and MATLAB (2016b, MathWorks). The following pre-processing pipeline was applied in all experiments: First, since we paused recording during the break between blocks, de-trending was applied using MATLAB, subtracting long-term linear trends from each block, thus zeroing the signal at beginning and end of blocks and avoiding discontinuities at the border between blocks. Then, further pre-processing was done in Analyzer, using the following pipeline: 0.1 Hz high-pass, zero-phase-shift 2nd order Butterworth filter; referencing to the nose electrode; correction of ocular artifacts using independent component analysis (ICA) (Jung et al., 2000) based on typical scalp topography and time course; and finally, discarding epochs that contained other artifacts. The latter part was semi-automatic – first, an algorithm marked artifacts based on predefined criteria: absolute difference between samples > 100 μV within segments of 100 ms; gradient > 50 μV/ms; absolute amplitude > 100 μV; or absolute amplitude < 0.5 μV for a duration of more than 100 ms. If an artifact was detected using any of these criteria, an epoch of 200 ms around it was marked. Then we performed visual inspection of all data to remove or add rare artifacts that were missed or marked by mistake. Artifact rejection was performed on 30 Hz low-passed data, causing the artifact rejection process to be blind to high frequency noise, which did not interfere with our analysis. Then, the data was exported from Analyzer to Matlab (prior to the 30 Hz low-pass filter). Finally, using MATLAB, a 1-20 Hz band-pass zero-phase-shift 4th order Butterworth filter was applied to the continuous data, followed by segmentation and averaging.

### Data processing

We calculated Event-Related Potentials (ERPs) locked to auditory stimulus presentation. The ERP amplitudes were measured from the midline central Cz electrode. This location was selected because it maximizes the N1 and P2 responses and is typically used to measure these components (e.g. as in Tremblay, Kraus, McGee, Ponton, & Otis, 2001). The data were parsed into segments beginning 100 ms before the onset of tone presentation and ending 400 ms after tone presentation. The average amplitude of the 100 ms pre-stimulus time served as a baseline for amplitude measurements. ERPs were obtained for each participant, by separately averaging trials of every block type and every tone, conditioned on every possible previous tone. This resulted in 5 tones x 4 previous tones (because there were no repetitions) x number of block types (2, 1 or 5 in Experiments 1, 2 and 3, respectively) ERPs. We then calculated the peak amplitudes of the N1 and P2 components for each ERP yielding 40, 20 or 100 measurements per participant in Experiments 1, 2 and 3 respectively.

N1 and P2 peak amplitudes were calculated in two stages. First, to minimize misidentification of peaks due to noise, we determined the peak time from the average of all presentations of each tone (i.e. not conditioned on the previous tone frequency) for each block type and participant. The resulting ERPs were based on a large number of trials and had satisfactory signal-to-noise ratio. We defined the N1 latency as the time of the absolute minimum (most negative) in the time window between 50 and 150 ms after stimulus onset and the P2 latency as the time of the absolute maximum in the time window between 130 and 250 ms after stimulus onset. If the peak latencies corresponded to the edge of the corresponding time windows, the participant was excluded from data analysis. In total, 0, 1 and 2 participants were excluded for this reason from Experiment 1, 2 and 3 respectively, leaving 79 participants overall in the analysis. We then computed the average voltage in a 12-ms window around the detected peak time for every possible combination of tone frequency with a previous tone frequency. Thus, peak latencies were determined for each tone frequency and block type regardless of previous tones, whereas the peak amplitudes were measured around these latencies, contingent on the previous tones. Overall this resulted in 4480 points for each potential type (840, 540 and 3100 for experiments 1, 2 and 3 respectively), or 8960 altogether.

### Statistical analysis of ERP peak potentials

To test the effect of both long- and short-term context on N1 and P2 peak amplitudes, we used linear mixed effect models (LME). LME were run in Matlab 2016b using the *fitlme* function. To be able to compare between the N1 and P2 amplitudes, the data were standardized using a z-score transform on the N1 and P2 separately, after multiplying N1 data points by −1. N1 and P2 amplitude values were modeled using 8 fixed factor predictors, 4 for each potential type. The 4 predictors were: an intercept, two continuous slope variables termed: *Interval-Mean* (long-term context: frequency interval between current tone and mean frequency overall in the sequence, semitones), *Interval-Previous* (short-term context: interval between current and previous tone frequencies, semitones) and another slope variable representing interaction between the two latter variables, encoded as their product: *Interval-Mean*Interval-Previous*. Random effect factors were added for all of the 8 fixed factors, grouped by participant number. In general, to determine whether a factor should be part of the model, we compared the two (nested) models trained with and without this specific factor using a likelihood ratio test (matlab *compare* routine). If the likelihoods of the two models were not significantly different that factor was excluded. Thus, in this model we omitted the random slope of the interaction variables *Interval-Mean*Interval-Previous* (both for N1 and P2) since they did not contribute to the overall explained variance (Likelihood ratio test between the model with and without these factors; χ^2^(2)=3, p=0.21). This resulted in 6 random effect terms in the LME. Thus, the LME model is described with the following Wilkinson formula (Wilkinson and Rogers, 1973):

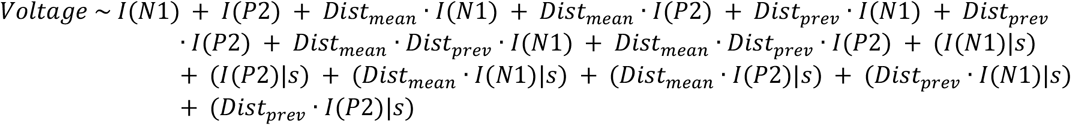

Where *Voltage* is either N1 or P2 standardized amplitudes of the responses to a specific tone (given all previous tone possibilities, for each experiment, condition and participant). *I*(*N*1) and *I*(*P*2) are indicator functions for N1 or P2 (each being 1 when the voltage belongs to the corresponding class and 0 otherwise). They therefore represent separate intercepts for N1 and for P2. *Dist_mean_* stands for *Interval-Mean, Dist_prev_* stands for *Interval-Previous*. For example, *Dist_prev_* · *I*(*N*1) denotes the *Dist_prev_* slope variable contributing to the N1 amplitudes. (X|s) denotes the random variable *X* grouped by participant number. The variable X is always assumed to be normally distributed with a mean of 0, and its variance is estimated from the data. Thus, (*I*(*N*1)|*s*) denotes a subject-specific contribution to the intercept for the N1 measurements; (*Dist_prev_* · *I*(*N*1)|*s*) is a subject-specific contribution to the corresponding slope. This LME model was estimated from data points from all 3 experiments together, resulting in 8960 data points overall (see end of *Data Processing* section above), collected from 79 participants.

Note that this is almost identical to modeling each potential type (N1 or P2) separately by:

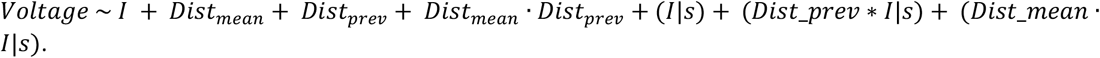

However, there are some distinctions. For instance, in the way we estimated the model the residual error is calculated overall from all data together and therefore it is more appropriate for statistical comparisons between the estimates of N1 and P2, which was one of our main goals.

For each fixed effect coefficient, a standardized effect size, Cohen’s d, was computed by dividing the estimate by its standard deviation (SD). The SD was calculated from the estimate standard error (SE) provided by fitlme, multiplying it by the square root of the number of degrees of freedom, DF (DF = 78; number of participants – 1). The significance value of each individual coefficient (ANOVA comparing it to 0) was given by the *fitlme* model output. To statistically compare between the contributions of two (or more) coefficients we ran a post-hoc coefficient test (F-test) for LME estimates, using the *coefTest* function in Matlab. Cohen’s d of these effects was calculated as the square root of F/DF (DF=78, as above).

To ensure the robustness of the results, we also ran a 2-level analysis, commonly used for example to analyze group results in functional MRI studies. A linear regression was run for each participant with regressors similar to the fixed effects above. We then performed a second-level analysis of the estimates using paired t-tests on participant-specific estimate values (see supplementary Figure S1 and Table S1).

To test the interaction of the overall frequency range of tones in the sequence with the effects estimated by the LME analysis described above, we ran another LME model adding interactions with the continuous slope variable *range* (overall frequency range in the sequence, semitones: Wide: 39, Medium: 28 and Narrow: 14). For each term in the above model we added another interaction term with *range*. This resulted in a large number of variables in the model and therefore we omitted the higher order interactions that did not contribute to the overall explained variance, according to a likelihood ratio test as described above. All random terms including the *range* variable as well as both fixed and random effects of the *Interval-Mean*Interval-Previous* (with and without *range* interaction) terms did not contribute to the overall explained variance (Likelihood-ratio test between the models with and without all of the latter terms; χ^2^(14)=24.9, p>0.99) and therefore were omitted from this model. However, since the fixed factors of the interaction terms *Interval-Mean*Interval-Previous* were included in the previous model, we estimated as well a model including these variables and verified that the results were comparable with and without them (supplementary Table S3). This LME model was run only on data points from Experiment 3, in which the range was manipulated within subject, resulting in 6200 data points (see end of first paragraph of *Data Processing* section above) collected from 31 participants. Effect sizes were calculated similar to the above, using DF=31-1=30.

### Single trial EEG amplitude extraction

We calculated single trial EEG amplitudes as the average voltage in a window of 12 ms centered around the latency of the N1 and P2 as determined by the subject’s ERP (i.e. average across trials). After excluding trials with electric artifacts, we were left with 2626, 1090, 7379 observations per participant on average in experiments 1, 2 and 3, respectively (313,366 observations overall were used to train the model). We also fitted the model separately for each of the frequency range conditions in Experiment 3. The total number of observations used in each of these conditions was 45,714, 91,979 and 91,072, respectively. (Recall that there were 2 versions of the Medium and Narrow conditions, Figure 1A, hence the larger number of trials).

### Adaptation Model

We used frequency-specific adaptation equations (Herrmann et al. 2013, 2014, 2015) to model single trial N1 and P2 responses. The model assumes frequency-tuned neural populations with Gaussian response profiles over a log-frequency axis. Each Population has resources, which determine the size of the responses that it can generate and are depleted in proportion to these responses. Therefore, resource depletion causes the contextual attenuation of the responses. Between sound presentations, resources recover exponentially with time. The amount of resources that are depleted is termed response adaptation (RA) following Herrmann et al. (2013, 2014, 2015). The RA values just before the presentation of a tone were calculated recursively for each specific stimulus sequence, trial by trial, using Equation 1.

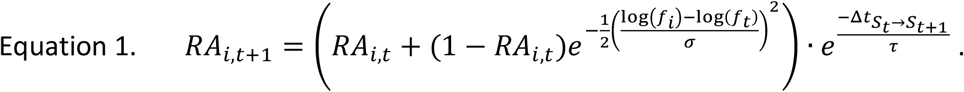

Here *RA_i,t_* is the response adaptation of neural population i centered around frequency *f_i_*, at time step t of the stimulus sequence, in which stimulus *S_t_* with frequency *f_t_* was presented. RA ranges between 0 (no adaptation) and 1 (maximal adaptation) and therefore 1-RA is the amount of available resources, and is therefore proportional to the response of neurons (full adaptation corresponds to minimum responsiveness and vice versa). The Gaussian term determines the amount of response evoked from population *i* by the stimulus presented at time t, and therefore of additional resource depletion that is determined by the interval between the current tone presented in the sequence and, the best frequency of neural population *i*. Δ*t*_*S*_*t*_→*S*_*t*+1__ is the time interval passing between the onset of the stimulus at time step t and the next stimulus at time step t+1. During this time, resources recover and thus RA decreases exponentially. There are two parameters in Equation 1: the Gaussian width of the frequency profiles - σ, and the time constant of the exponential recovery - *τ*. These were termed together 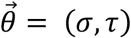. Once 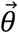 is given, Eq. 1 allows the computation of the adaptation level for each neuronal population and at each time point.

Next, the model assumes a linear relationship between the measured EEG data and RA of the population at the presented tone frequency (Equation 2).

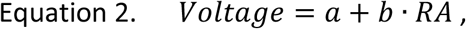

where *Voltage* is the measured EEG amplitude, either N1 or P2, and *a* and *b* are the linear factors associating RA and the data. Since RA is defined between 0 and 1, these factors allowed for appropriate scaling and shift to the EEG units of measurement. The inverse relation between RA and the responsiveness of neurons (1-RA) is captured by the values of *a* and *b*.

*RA_i,t_* was first computed for all the 5 tone frequencies i and all t time steps in a specific sequence. Each participant heard different stimulus sequences and therefore RA was computed for each participant separately. Then, to predict responses at each time step t, the relevant RA was taken as that of the i corresponding to the currently presented tone, as in Herrmann et al. (2013, 2014, 2015). The last step assumes that EEG responses measured at time step t are dominated by neural population centered around the frequency of the presented stimulus *f_i_* = *f_t_*.

The procedure we used was the one used by Hermann et al. (2013, 2014, 2015), and we used it in order to be consistent with them. A more natural procedure would consist of integrating the RA values for all neuronal populations, weighted by their tuning profiles (the exponential term in Eq. 1). We verified that the two methods are comparable by calculating the full RA predictions, weighted by the tuning profiles, for one specific participant and plotting them against the simplified model. We show that the two models are strongly linearly dependent (supplementary Figure S3).

### Model fit and parameter estimation

All data analysis related to model fitting was carried out using Matlab 2016b (Mathworks, MA, USA). In total, the model had four free parameters – The two mechanistic parameters 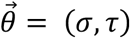, and the linear factors *a* and *b*. All parameters were estimated from the data. However, 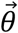 and the linear factors were treated differently. 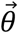 are the parameters of interest for this study, while the linear factors are ‘nuisance’ parameters that have to be fitted but are not interpreted. Therefore, fitting the model was done in two steps. First, single-trial model predictions (RA) were calculated for a range of pre-defined possible 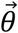 values; 18 values for *σ* spanning 1 to 18 semitones, and 25 for *τ*, spanning 0.2 to 5 seconds with 0.2 second steps (resulting in 450 possible 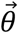 combinations). Second, for each 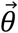, single trial EEG responses were regressed on the computed RA values. In practice, a linear mixed-effects model (LME) was fitted (using Matlab *fitlme*) using RA as a continuous fixed effect with random intercept and slopes grouped by participant (Wilkinson formula (Wilkinson and Rogers, 1973):

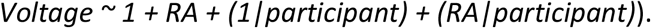

This approach made it possible to estimate participant-specific linear factors while reducing the amount of overfitting that would be generated by estimating the linear factors of each participant separately. Log-Likelihood (LL) statistics of the LME model were extracted for each value of 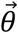. Then, 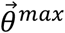, the 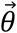 value resulting in the maximum likelihood, was selected as the best estimate for the parameters of interest.

### Testing significance of model fit

To test whether the adaptation model generally described the data better than chance we repeated all the steps of parameter estimation for randomly permuted measurements. For permuted data the model was not expected to perform better than chance and therefore this allowed calculation of the distribution of the LL statistic under the null hypothesis of no effect of sequence order. We used 250 permutations. In particular, for each permutation of the data, we collected the maximum log-likelihood values (over of all possible 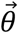 values), and plotted the null distributions of maximum LL values for N1 and P2 separately. The maximum LL of real data were compared to the null distributions of maximum LL.

### Confidence regions for the parameter estimates

To calculate a confidence interval around 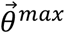, we asked which 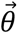 values are significantly different than 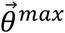. Due to Wilk’s theorem the quantity:

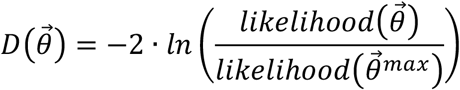

should be distributed as *χ*^2^ with 2 DF under the null hypothesis that 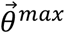 does not describe the data better than any other parameter value, and for a big enough sample size. We also verified this assumption empirically, simulating the null distribution of D and showing that it is comparable to *χ*^2^ with 2 DF at 2 representative values of 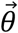 for N1 and P2 data (supplementary Figure S4). We thus calculated the D statistic for all possible 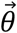 values and asked whether it is significant using a *χ*^2^ test with 2 DF. A large D is expected for 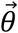 values that are significantly different from 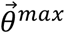. The statistical question of which 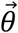 values are significantly different from 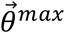 is equivalent to calculating a confidence region for 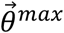. The D values at 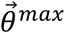 always equal to 0 by definition. The threshold of D<6, which is approximately the value corresponding to a p=0.05 of the *χ*^2^ distribution with 2 DF, was used to define the 95% confidence region around 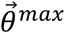.

### Comparing parameter estimates for N1 and P2

We statistically contrasted the values of 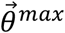 estimated for N1 vs. P2 using three methods. First, we compared the 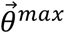 values of N1 and of P2 by checking whether the 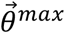 of, e.g. N1, fell outside the confidence region of P2 and vice versa (see previous section). Second, we performed bootstrap (random sampling with replacement) on the group of 79 participants (using all experiments together), and repeated the estimation procedure 100 times. This resulted in an estimate of the distribution of 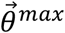 values (under the assumption that the group of subjects represents the population). The distribution of differences of 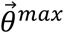 values calculated in the same bootstrap repetition for N1 vs. P2 was compared to 0. Third, we used data permutations to create the null distribution of the difference between LL at 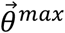 of the two potential types (termed log-likelihood differences, LLD). For each of 250 permutation repetitions, we flipped between N1 and P2 of single trials, with probability 0.5 per each trial flip. For each iteration, we repeated the estimation procedure and computed the LLD. We thus estimated the null distribution of LLD under the assumption that the parameters of N1 and P2 were identical, and calculated the p-value of the LLD of the actual data by comparing it to this null distribution.

### Comparing parameter estimates for different frequency ranges

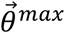 values were estimated separately for each of the stimulus frequency range conditions (Wide, Medium and Small) in Experiment 3. Parameter estimation was repeated 100 times using a bootstrap procedure – simulating new groups of participants by random sampling with replacement from the pool of 31 participants. To statistically test the effect of frequency range on adaptation bandwidth of N1 and P2 we fitted an LME model to the bootstrapped *σ^max^* values, using the Matlab *fitlme* function. The bootstrap repetition number was used as the grouping variable for the random effects. The stimulus frequency range variable was modeled as a continuous fixed effect with random intercept and slope. Thus, the LME model is described with the following Wilkinson formula:

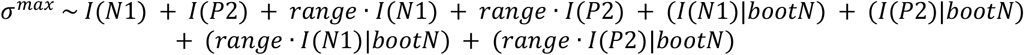

Where *σ^max^* are the estimated adaptation bandwidths in each of 100 bootstrap runs, in each of the 3 range conditions and both for N1 and P2 data (600 values in total), and *I(N1)* and *I(P2)* are indicator functions for N1 or P2, respectively. Thus, the *I* represent separate intercepts for N1 and P2. *Range* is the overall range of frequencies presented in a sequence, in semitones (so *range · I*(*N*1) represents the effect of range on the N1 bandwidths) and *bootN* is the bootstrap run number serving as a grouping variable for the random effects. Effect size (Cohen’s d) was calculated as above (DF=100-1=99, for 100 bootstrap repetitions).

## RESULTS

In 3 EEG experiments, 79 participants (21, 27 and 31 in Experiments 1, 2 and 3) were presented with sequences of 5 equiprobable pure tones (Figure 1A), which they were instructed to ignore while concentrating on a silent film. Using a passive paradigm allowed us to investigate neural effects that are elicited automatically and do not depend directly on attention or on any active task. In Experiments 1 and 2 we analyzed the control conditions from a previously published study (Regev et al., 2019) (the published study included also conditions involving rare deviants, which were the focus of that study; We do not analyze these conditions here as we focus on equiprobable sequences). The sequences from Experiment 1 and 2 we report here had a relatively wide frequency range (32 and 34 semitones between lowest and highest tones used in the sequence, respectively, Figure 1B, Methods). Experiment 3 included 3 frequency range conditions: Wide, Medium and Narrow (39, 28, and 14 semitones respectively). We examined the amplitudes of the N1 and P2 auditory-evoked responses and their dependence on two features of the tones along the sequence: 1) The frequency interval between the current tone and the mean sequence frequency (*Interval-Mean*), and 2) the frequency interval between the current tone and the previous tone frequency (*Interval-Previous*). *Interval-mean* represented a long-term context variable, because in order to show sensitivity to the overall mean in the sequence, integration of several previous tones should occur (at least 5 previous tones, ~2.5 s). *Interval-previous* represented a short-term context variable, at the scale of 1 SOA (~0.5 s).

### N1 but not P2 is sensitive to long-term context

Absolute N1 amplitudes increased as a function of the frequency interval between the current tone and overall mean frequency in the sequence. This dependence manifested itself as a typical inverted U-shape pattern, so that the most negative N1 amplitudes were elicited in response to the most extreme tones and the least negative N1 was elicited by the middle tone (which was also approximately equal to the mean frequency of the sequence). This phenomenon was robust and replicated in all 3 experiments (Figure 1C, E). In contrast, P2 amplitudes did not show significant dependence on the mean sequence frequency (Figure 1D, E). To quantify this effect, we used a linear mixed effect model (LME) including data from all experiments together (Table 1, Figure 2). The slope of the N1 amplitudes on *Interval-Mean* was significantly different from 0 (F(1,8952)=70, p=6.7E-17, d=0.94, Table 1), while the slope of the P2 amplitudes on *Interval-Mean* was not significantly different from 0 (F(1,8952)=0.15, p=0.7, d=-0.04). The difference between the N1 and P2 slopes for *Interval-Mean* was significantly different from 0 (F(1,8952)=35.5, p=2.6E-09, d=0.67, Table 1, Figure 2 – contrast #3). A two-stage procedure including linear regression on individual participants followed by second-level analysis at the group level gave similar results (supplementary Figure S1 and Table S1).

**Table 1.** Linear mixed effects (LME) results – effect of long- and short-term context on N1 and P2. **Top** – N1 and P2 amplitudes (standardized using a z-score transform after negating N1 data points) were modeled using the 8 predictors listed in the first (left-most) column. The model consisted of fixed and random factors (grouped by participant) for each of the listed predictors (except for the interaction term Interval-Mean*Interval-Previous which did not have a random factor since the latter did not contribute to explained variance), see Methods. Columns 2 to 5: Fixed-effect estimates (Estimate), standard errors of the estimates (SE), F-statistic used for ANOVA comparing the estimates to 0, with degrees of freedom, significance level (p-value) of the latter test, standardized effect size (Cohen’s d, Methods). The predictors Interval-Mean and Interval-Previous stand for the frequency interval between the current tone and sequence mean or current tone and previous tone frequency, respectively, in semitones. **Bottom** – Post-hoc pairwise coefficient comparison between predictors. The LME model was run on 8960 observations collected from 79 participants overall in the 3 experiments (Methods).

| Predictors |  | Estimate | SE | F(1,8952) | p-value | d |
| --- | --- | --- | --- | --- | --- | --- |
| Intercept | N1 | -0.51 | 0.078 | 42.2 | 8.8E-11 | -0.73 |
|  | P2 | -0.18 | 0.074 | 5.80 | 0.016 | -0.27 |
| Interval-Mean | N1 | 0.04 | 0.004 | 70.0 | 6.7E-17 | 0.94 |
|  | P2 | -0.002 | 0.005 | 0.15 | 0.7 | -0.04 |
| Interval-Previous | N1 | 0.01 | 0.003 | 26.9 | 2.2E-7 | 0.58 |
|  | P2 | 0.02 | 0.003 | 31.8 | 1.7E-8 | 0.63 |
| Interval-Mean *<br>Interval-Previous | N1 | -0.0008 | 0.0002 | 10.08 | 0.0015 | -0.36 |
|  | P2 | -0.0002 | 0.0002 | 0.96 | 0.32 | -0.11 |

| Pairwise comparisons between predictors | F(1,8952) | p-value | d |
| --- | --- | --- | --- |
| N1 vs. P2 Interval-Mean | 35.5 | 2.6E-9 | 0.67 |
| N1 vs. P2 Interval-Previous | 0.23 | 0.62 | -0.05 |
| N1 Interval-Mean vs. -previous | 25.4 | 4.6E-7 | 0.57 |
| P2 Interval-Mean vs. -previous | 21.41 | 3.7E-6 | -0.52 |
| N1 Interval-Mean vs. -previous<br>vs.<br>P2 Interval-Mean vs. -previous | 46.3 | 1.03E-11 | 0.77 |

**Figure 2.**
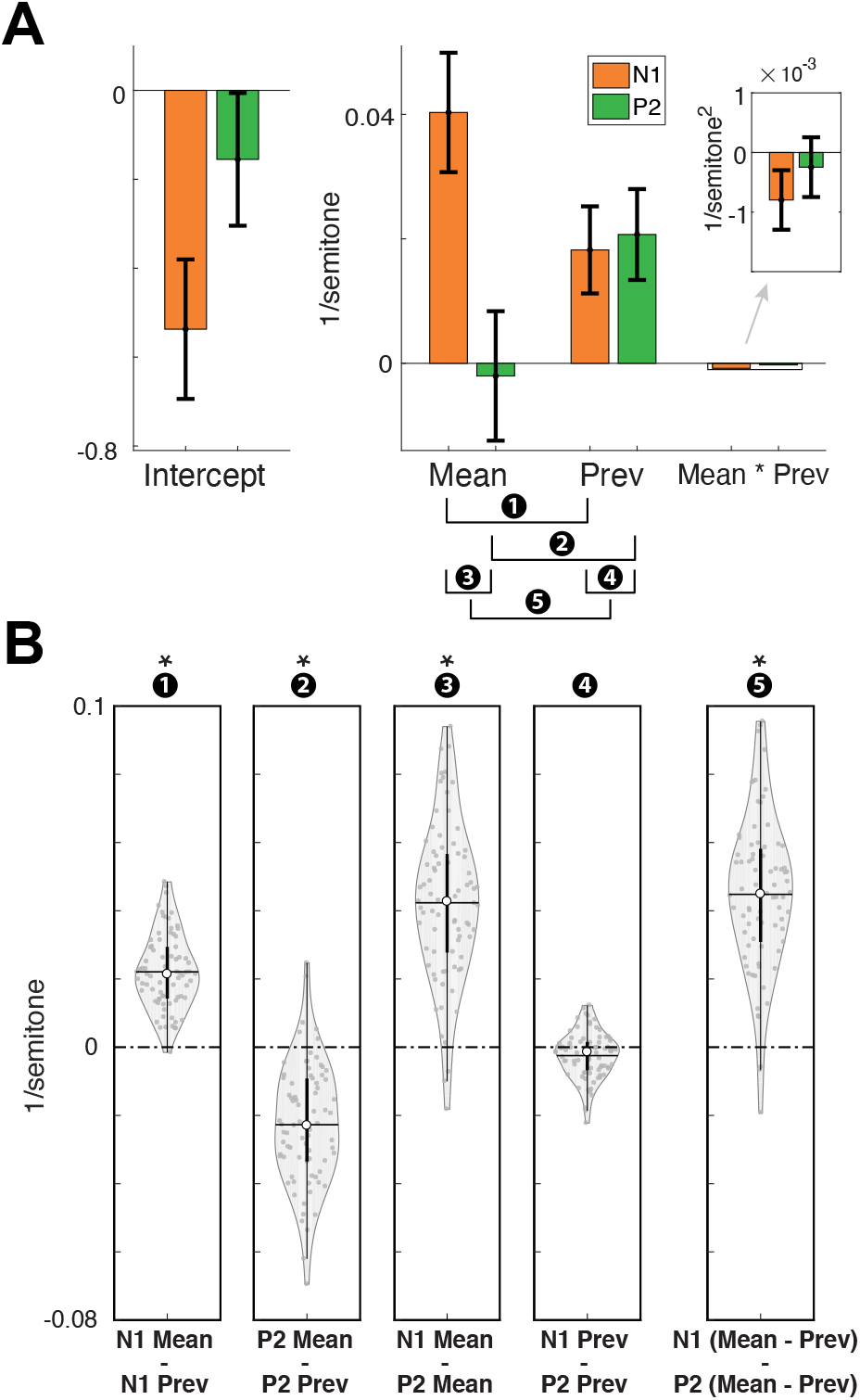
long- and short-term context effects on N1 and P2 amplitudes. **A** – Bar-graphs illustrate fixed-effect estimates values from a linear-mixed effects (LME) model (Table 1 and Methods for further specification). ‘Mean’ and ‘Prev’ stand for Interval-Mean and Interval-Previous, denoting the frequency intervals between the current tone and the sequence mean or previous tone, respectively, in semitones. The predicted N1 and P2 voltages were z-scored (after reversing the sign of the N1 data points). Error-bars represent 95% confidence intervals around the estimate (calculated by multiplying the SE of the estimate by the 95% inverse t-distribution value (DF=78)). **B** – Violin plots illustrate comparisons between LME estimates. Each dot represents one participant. White numbers in black circles above the violin plots indicate to which comparison they correspond (displayed under A). The significant contrasts are marked with an asterisk. Participant-specific estimates were calculated by adding the common fixed-effect estimates to participant-specific random effects. Horizontal lines represent the mean, white circles the medians, and thick and thin black vertical lines represent the 25% and 75% percentiles, respectively.

### P2 is more sensitive than N1 to short-term context

N1 and P2 absolute amplitudes increased as a function of the interval between the current and previous tone frequencies, but this effect was larger for P2 than for N1 (Figure 3). To visualize this, we pooled the possible combinations of current and previous tones according to the ‘degree of neighborhood’. Neighbor 1-4 denotes the proximity of tones on the frequency axis in a specific sequence (Figure 3A, B). Figure 3 (C, D) compares the N1 and P2 peak amplitudes for when the previous stimulus was ‘Neighbor 1’ vs. ‘Neighbor 2’. We concentrated just on the ‘Neighbor 1’ and ‘Neighbor 2’ groups since they were comprised of more combinations of current and previous tones and included all current tones, whereas by design only extreme tones in every sequence could have neighbor ‘3’ and ‘4’. The difference between ‘Neighbor 1’ and ‘Neighbor 2’ amplitudes was larger for P2 than for N1 in 7 out of 8 block types overall in the 3 experiments (Figure 3D).

**Figure 3.**
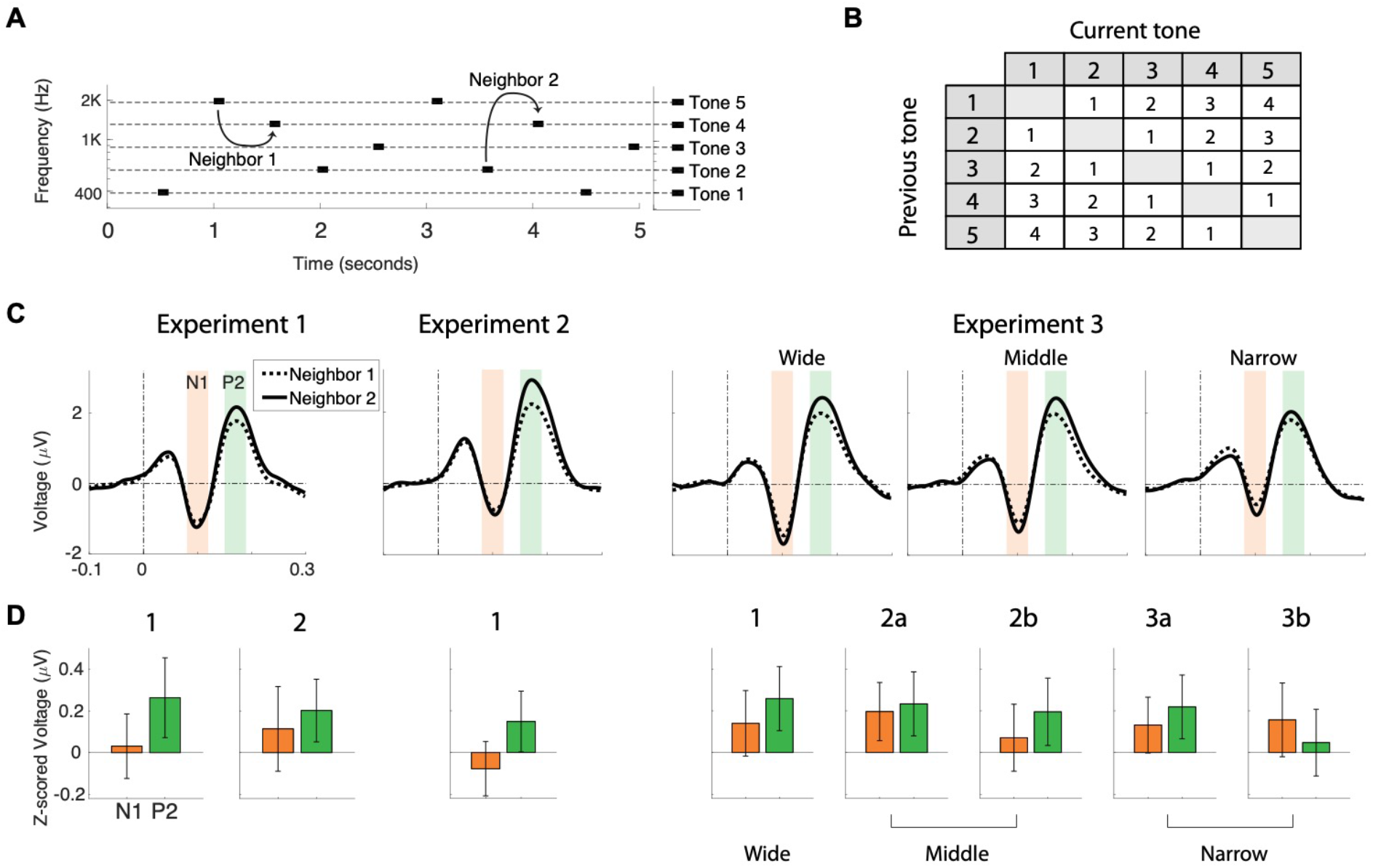
P2 is more sensitive than N1 to short-term context. **A** – Example stimulus sequence denoting one frequency interval between two consecutive tones that are 1^st^ order neighbors on the sequence frequency axis (‘Neighbor 1’) and the 2^nd^ order neighbors (‘Neighbor 2’). **B** – All possible combinations of current and previous tones in a sequence, and their grouping into the ‘degree of neighborhood’. Column and row headers of the table denote ordinal tone numbers (1 to 5 from low to high frequencies, see right axis in A). Numbers inside the table denote the ‘degree of neighborhood’. **C** – ERPs of ‘current tones’ when the previous tone was ‘Neighbor 1’ and ‘Neighbor 2’ for each experiment and condition separately. Shaded orange and green areas illustrate the time windows across which the N1 or P2 voltages were averaged. **D** – Bar-graphs denote the mean and 95% confidence intervals (across participants) of the difference between peak amplitudes in the ‘Neighbor 2’ and 1’ conditions. Peak amplitudes were z-scored for N1 and P2 separately (after reversing the sign of the N1 data points).

The LME model including data from all experiments together (Table 1, Figure 2) indicated that the frequency interval between the current and previous tone (*Interval-Previous*, semitones) significantly affected both N1 (ANOVA comparing the LME estimates to 0; F(1,8952)=27, p=2.2E-7, d=0.59, Table 1) and P2 (F(1,8952)=31.8, p=1.7E-8, d=0.64) amplitudes. The effect size of *Interval-Previous* was nominally larger for P2 than for N1 but they were not significantly different (F(1,8952)=0.24, p=0.62, d=-0.05). Regressions on individual participants and second-level analysis of regression estimates gave similar results (supplementary Figure S1 and Table S1). Notably, excluding the interaction term between *Interval-Mean* and *Interval-Previous* from the LME model reduced the short-term (*Interval-Previous*) context effect for N1 but not for P2 (Supplementary Table S2), resulting in a significant difference between N1 and P2 *Interval-Previous* effect (F(1,8954)=9, p=0.0025, d=-0.34). Thus, whereas the dependence of P2 on short-term context was robust, the dependence of N1 on short-term context interacted with its dependence on long-term context.

### Sequence frequency range affects N1 but not P2 amplitudes

N1 amplitudes were reduced when the frequency range in the sequence was smaller, while P2 amplitudes were not affected much by the frequency range manipulation (Figure 1C and D, Experiment 3), consistent with the fact that N1 amplitudes were more affected by long-term adaptation throughout the sequence than P2. To test this statistically we ran another LME model including *range* as a predictor (Table 2). This analysis confirmed a significant contribution of *range* to the N1 intercept (ANOVA comparing LME estimates to 0; F(1,6188)=36, p=1.9E-9, d=1.1) but not to the P2 intercept (F(1,6188)=1.8, p=0.18, d=0.24) and a significant difference between the effect of range on N1 and P2 intercepts (F(1,6188)=10.9, p=9.5E-4, d=−0.6).

**Table 2.**
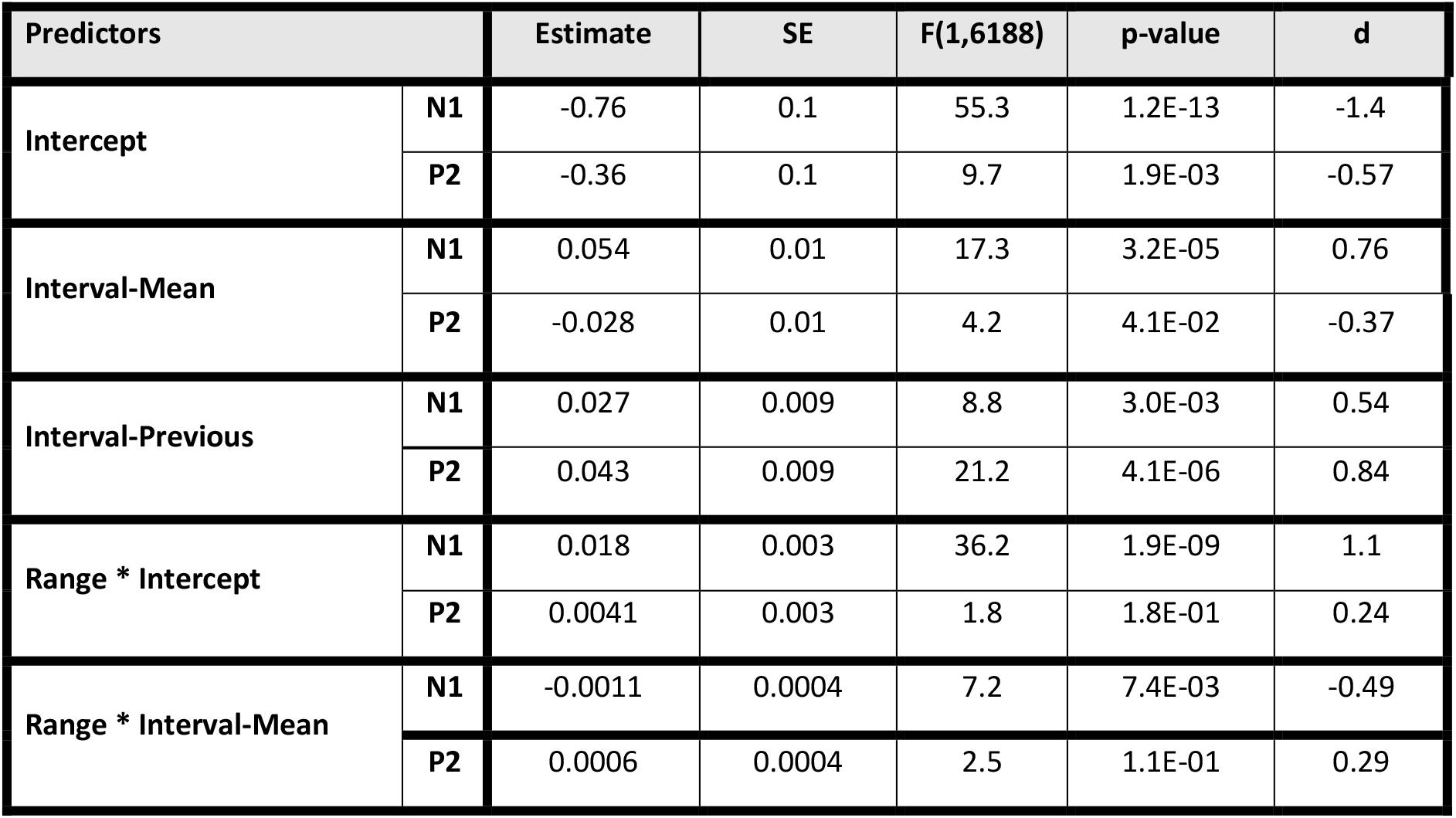
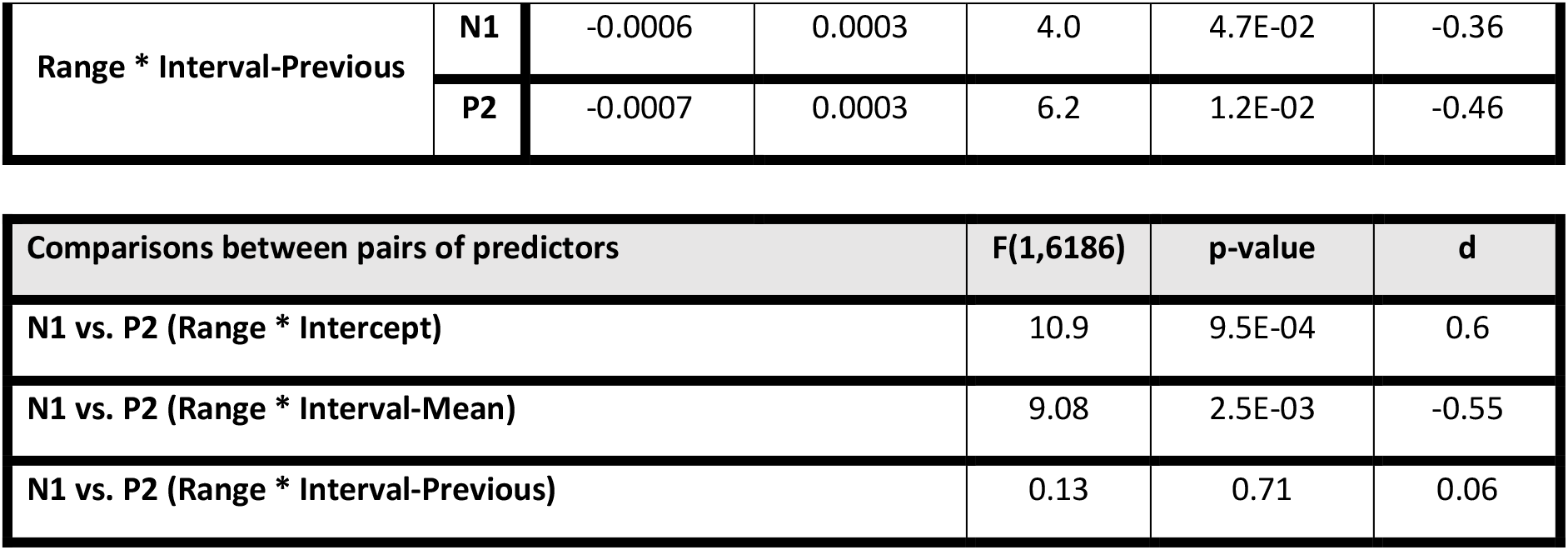
Linear mixed effects (LME) results including interactions with frequency range. Entries are similar to Table 1. Here only data from Experiment 3 (31 participants) was used to train the model, resulting in 6200 observations for N1 and P2 altogether (Methods).

| Predictors |  | Estimate | SE | F(1,6188) | p-value | d |
| --- | --- | --- | --- | --- | --- | --- |
| Intercept | N1 | -0.76 | 0.1 | 55.3 | 1.2E-13 | -1.4 |
|  | P2 | -0.36 | 0.1 | 9.7 | 1.9E-03 | -0.57 |
| Interval-Mean | N1 | 0.054 | 0.01 | 17.3 | 3.2E-05 | 0.76 |
|  | P2 | -0.028 | 0.01 | 4.2 | 4.1E-02 | -0.37 |
| Interval-Previous | N1 | 0.027 | 0.009 | 8.8 | 3.0E-03 | 0.54 |
|  | P2 | 0.043 | 0.009 | 21.2 | 4.1E-06 | 0.84 |
| Range * Intercept | N1 | 0.018 | 0.003 | 36.2 | 1.9E-09 | 1.1 |
|  | P2 | 0.0041 | 0.003 | 1.8 | 1.8E-01 | 0.24 |
| Range * Interval-Mean | N1 | -0.0011 | 0.0004 | 7.2 | 7.4E-03 | -0.49 |
|  | P2 | 0.0006 | 0.0004 | 2.5 | 1.1E-01 | 0.29 |
| Range * Interval-Previous | N1 | -0.0006 | 0.0003 | 4.0 | 4.7E-02 | -0.36 |
|  | P2 | -0.0007 | 0.0003 | 6.2 | 1.2E-02 | -0.46 |

| Comparisons between pairs of predictors | F(1,6186) | p-value | d |
| --- | --- | --- | --- |
| N1 vs. P2 (Range * Intercept) | 10.9 | 9.5E-04 | 0.6 |
| N1 vs. P2 (Range * Interval-Mean) | 9.08 | 2.5E-03 | -0.55 |
| N1 vs. P2 (Range * Interval-Previous) | 0.13 | 0.71 | 0.06 |

### Sequence frequency range attenuates long- and short-term context effect

The latter LME model also indicated that both the short- and long-term context effects were attenuated for sequences with larger frequency ranges, consistent with adaptation with a limited bandwidth. There was a significant interaction between the range and the short-term context variable *Interval-Previous*, such that for both N1 and P2, the effect of *Interval-Previous* was smaller the larger the range was (N1: F(1,6188)=4, p=4.7E-2, d=-0.36), P2: F(1,6188)=6.2, p=1.2E-2, d=-0.46). The interaction between range and the long-term context variable *Interval-Mean* was significant for N1 (F(1,6188)=7.2, p=7.4E-3, d=-0.49, smaller effect of *Interval-Mean* with larger range; Table 2) but not for P2 (F(1,6188)=2.5 p=0.1, d=0.3). Notably, the interaction terms *interval_mean*interval_previous* did not contribute significantly to this model so we omitted them (Methods), however see *Supplementary Table S3* for comparison to a model including these terms, which gave similar results.

In summary, the ERP results demonstrated that N1 and P2 were affected differently by context: N1 was highly affected by long-term context (*Interval-Mean*) and P2 was not. Additionally, both were affected by short-term context (*Interval-Previous*) but P2 more robustly so. Furthermore, the spectral context (frequency range in the sequence) had a distinct effect on the N1 and P2 amplitudes: smaller sequence range reduced N1 more than P2 amplitudes, suggesting that adaptation affects the N1 more than it affects the P2. These results imply that N1 and P2 have distinct time scales of contextual influences. Additionally, the effects of long- and short-term context were generally reduced for larger frequency ranges, suggesting that some limited frequency bandwidth plays a role in these effects.

The specific values of the time scales and frequency bandwidth cannot be directly computed using the ERP analysis presented until here. Importantly, the two context predictors we used in the LMEs; *Interval-Mean* and *Interval-Previous* made it possible to consider only very short-(1 previous tone) or very long-term (the sequence mean) contextual effects. In order to estimate the relevant temporal and spectral scales of these contextual effects, we employed a computational model.

### Adaptation model

We hypothesized that both the N1 and P2 results could be generated by a single underlying neural mechanism – adaptation of frequency-selective neural populations with relatively wide bandwidths that have two different time constants.

In the auditory system, frequency-selective neurons respond not only to their best frequency but also to nearby frequencies. Therefore, presenting a tone would adapt not only neural populations tuned exactly to that tone’s frequency but also populations tuned to nearby frequencies. Further, if the interval to the next tone is short enough relative to the time scale of adaptation recovery, this adaptation would not recover fully before the next stimulus occurs. Thus, given that effective frequency response profiles of neuronal populations are wider than the frequency intervals between tones in a stimulus sequence, cross-frequency adaptation (Taaseh, Yaron, & Nelken, 2011; also termed co-adaptation, Herrmann et al., 2015; Herrmann, Schlichting, & Obleser, 2014) would render the adapting populations sensitive to frequency intervals. Moreover, the time it takes for neurons to recover from adaptation determines the duration of this effect. If recovery rates are slow relative to the inter-stimulus interval, neurons would accumulate adaptation due to their responses to more than one previous tone in the sequence.

With these premises, we used computational modelling (see Methods for equations) to test the feasibility of adaptation as the neural mechanism accounting for the ERP results presented above, and to estimate quantitatively the effective time and frequency scales underlying the context-sensitivity of the N1 and P2 potentials. A similar modelling approach was applied in the past for neural responses in rats (Taaseh et al., 2011). Further, this model was applied for EEG by Herrmann et al. (2013, 2014, 2015) and we used a similar formulation to the latter studies for comparability. We fitted model predictions to single trial N1 and P2 amplitudes and estimated *σ*, the bandwidth of frequency response-adaptation profiles, and *τ*, the time constant of recovery from adaptation (Figure 4), for N1 and P2 separately.

**Figure 4.**
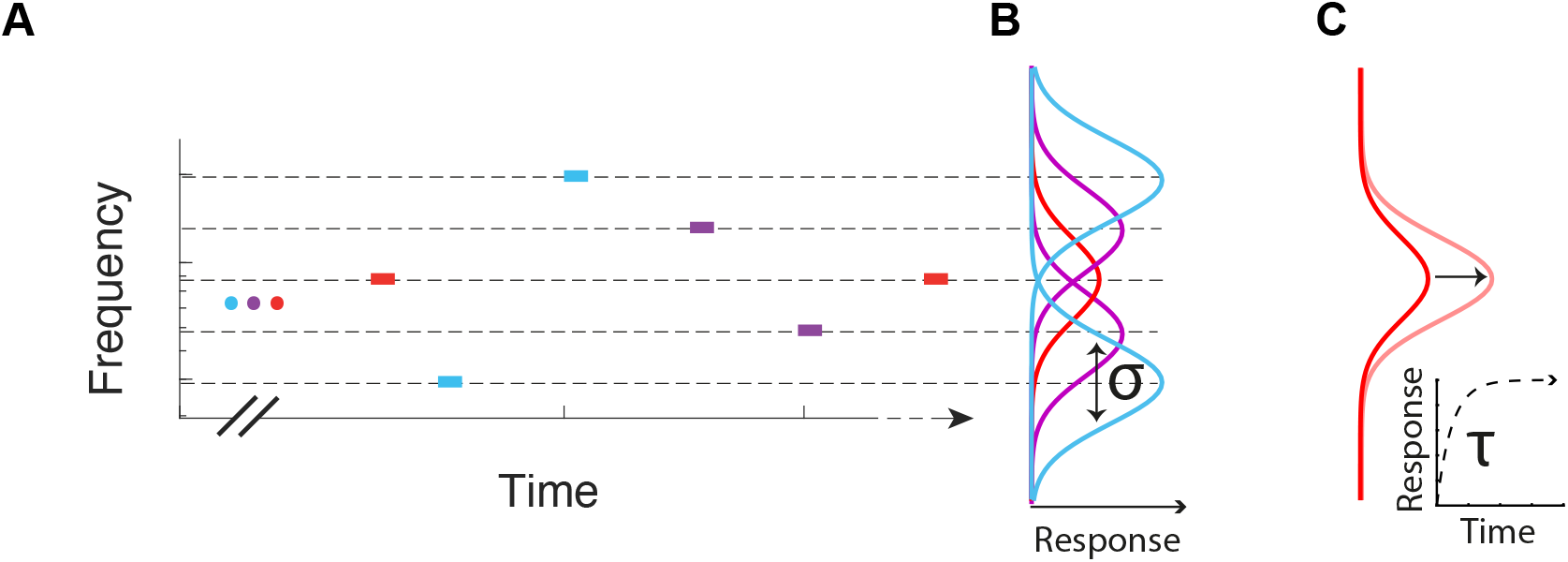
Illustration of the adaptation model. **A** – An example segment of a tone sequence in the experiment serving as stimulus. Color code as in Figure 1. **B** - Schematic Gaussian frequency response-adaptation curves of neural populations assumed by the model. All curves have the same bandwidth σ and each curve is centered around 1 of the 5 tone frequencies in the sequence. The color of the curves matches the color of the stimulus (A) at its best frequency. The amplitudes of the curves represent the average adaptation of the neural populations throughout the sequence (excluding 5 initial tone presentations). Populations with response-adaptation curves centered around the middle frequency (red), respond most frequently throughout the sequence and therefore have the most adapted (attenuated) response-adaptation profile. **C** - Exponential recovery from adaptation. The red curves represent the frequency response-adaptation curves of a population at two time points during a period with no tone presentations. Inset below is an exponential recovery function for a given time constant τ. The curve increases along the arrow connecting the dark to the light red curves, as specified by the exponential function in the inset.

### The adaptation model accounts for both N1 and P2 data

The model was fitted to single trial N1 and P2 data separately and the values of the time and frequency scale parameters, 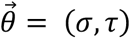, were estimated by selecting the 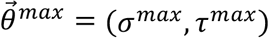 values maximizing the log-likelihood (*LL^max^*), using a search over a predetermined grid of parameter values. To test the significance of the fit, we fitted the model to surrogate data consisting of random permutations of the measured responses across time (Methods). For both N1 and P2 the *LL^max^* values obtained using the actual data were much larger than all null *LL^max^* values obtained from the surrogate data. This indicated that the adaptation model fitted the data better than chance (p<0.004, since real *LL^max^* for both N1 and P2 was larger than all *LL^max^* calculated in 250 repetitions, supplementary Figure S5).

### N1 has a longer adaptation recovery time than P2

The estimated time constant for recovery from adaptation, *τ^max^*, was consistently longer for N1 relative to P2. This result was found when fitting the model using data from Experiments 1, 2 and 3 separately, as well as when using the data from all experiments together. The values of *τ^max^* were 5, 4.6, 2.4 or 3.2 s for N1 and 0.4, 0.8, 1 or 1 s for P2 in Experiments 1, 2, 3 or when using all data together, respectively (Figure 5A). In Experiment 1, *τ^max^* of N1 was on the upper boundary of the allowed parameter range (5 s, equaling 2 repetitions of a 5-stimulus sequence). We limited the *τ* scale to 5 s since the predicted values become almost constant for larger *τ* values, due to the fact that the stimulus sequence was composed of successive permutations of the five frequencies. Therefore, a time constant of 5 s should be interpreted as 5 or longer.

**Figure 5.**
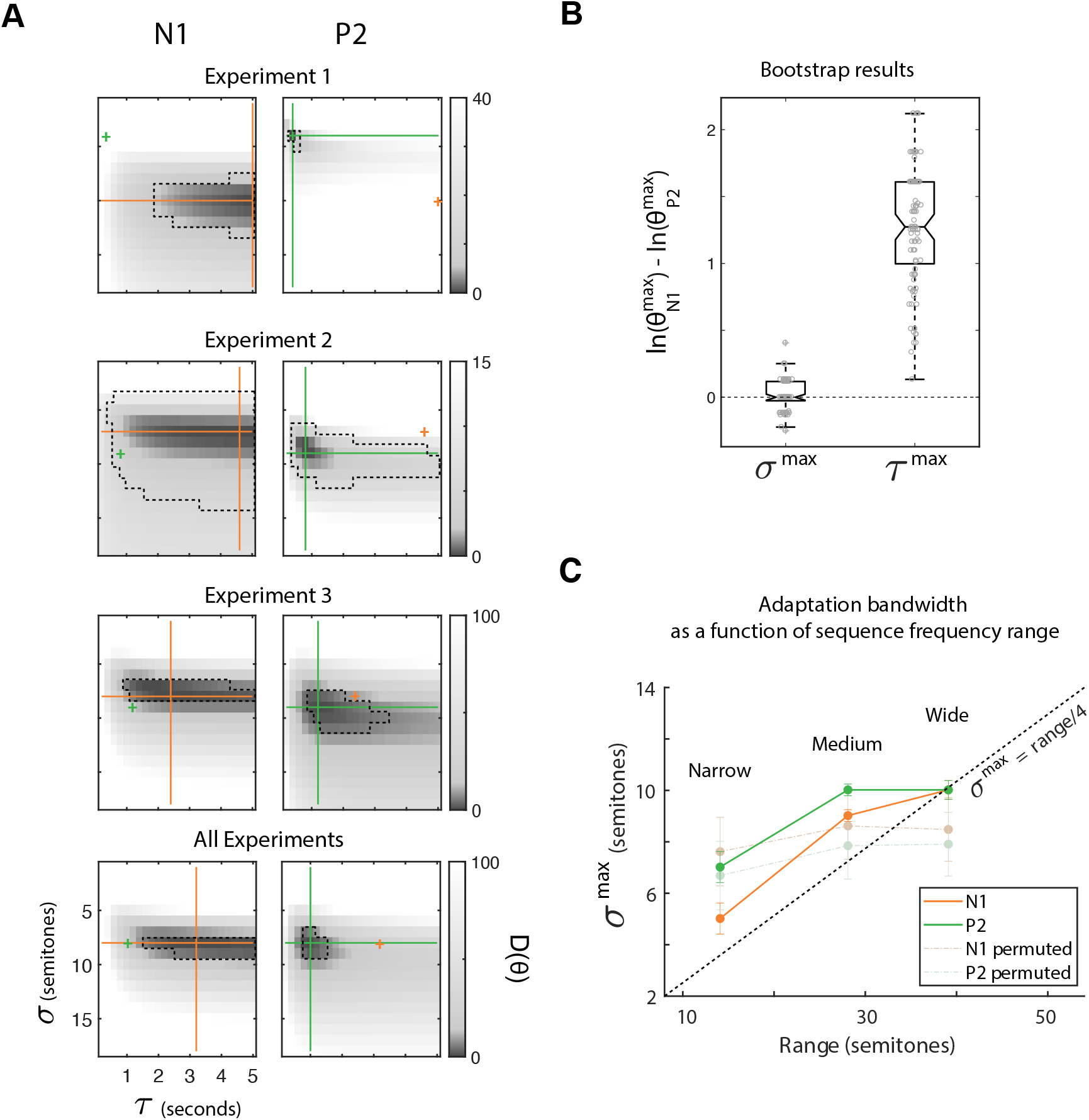
Adaptation model reveals the time and frequency scales of N1 and P2 context sensitivity. **A** - D values (−2* log-likelihood ratio relative to 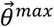 which maximizes the likelihood, Methods), for each possible value of 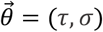, for each experiment, for N1 (left) and P2 (right). Cross-hairs (N1: orange, P2: green) are located at the maximum-likelihood estimated parameter values (i.e. at 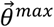). Small cross-signs are located at 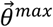 of the other potential type (exactly the crossing point of the neighboring plot, same color code), for visual comparison between 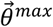 of the two potential types. Dashed lines surround 95% confidence regions for 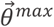 calculated according to the null distribution of D (χ^2^ with 2 df, Methods). **B** - Bootstrap results comparing estimated parameter values for N1 and P2. Each gray dot is the difference between 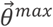 estimated for N1 and for P2 in one of 100 bootstrap repetitions (sampling with replacement from the 79 participants and repeating the full parameter estimation procedure, Methods). **C** – Frequency bandwidth depends on spectral context. σ^max^ as a function of stimulus frequency range for N1 (orange), P2 (green) and for N1 and P2 after permuting the order of trials within each range condition 250 times (dashed, see legend). Data from Experiment 3 only. Error bars of the N1 and P2 plots are 95% confidence intervals estimated from 100 bootstrap repetitions sampling with replacement the 31 participants and repeating parameter estimation procedure. Error bars for the permuted data plots are 95% confidence intervals calculated from the 250 permutations.

The difference between *τ^max^* of N1 and of P2 was significant. We used three methods for comparing them (Methods): (1) The values of *τ^max^* of each potential type fell outside the 95% confidence region of *τ^max^* of the other type (Figure 5A); (2) *τ^max^* of N1 was larger than *τ^max^* of P2 in all bootstrap repetitions we performed (sampling with replacement over the 79 participants and repeating the parameter estimation procedure; Figure 5B, p<0.01 since 100 bootstrap repetitions were conducted); and (3) the difference between the log-likelihoods calculated at *τ^max^* and at *τ^max^*of the other potential type, were significantly larger than their null distribution, estimated by fitting the model to permuted data (p=0.0039 for N1 and p=0.012 for P2, supplementary Figure S6).

### N1 and P2 have similar frequency bandwidths of adaptation

In contrast to the time constant, the estimated frequency bandwidth, *σ^max^*, was similar for N1 and for P2. *σ^max^* was 10, 7, 8 or 8 semitones for N1 and 4, 9, 9, or 8 semitones for P2 in Experiments 1, 2, 3 or all together, respectively (Figure 5A). In fact, using the data of all experiments together, *σ^max^* was exactly 8 semitones for both N1 and P2 (Figure 5A lowest panels). Additionally, the differences between *σ^max^* values of N1 and of P2 estimated for 100 bootstrap repetitions of the 79 participants were not significantly different than 0 (Figure 5B).

### Adaptation bandwidth rescales to the sequence frequency range

Next, we asked whether the parameters of the model (*σ^max^* and *τ^max^*) are constant when the spectral context changes. We fitted the model and estimated parameter values separately for each of the frequency range conditions in Experiment 3. First, we tested whether allowing the time constant, *τ^max^*, to vary for different ranges improves significantly the model fit. Estimating a separate *τ* did not increase significantly the explained variance neither for N1 (Likelihood ratio test comparing models with constant or varying *τ*: *χ*^2^=0, df=2, p=1) nor for P2 (*χ*^2^=4.7, df=2, p=0.09). We therefore estimated a single time constant for all the possible ranges. Next, we tested whether the frequency adaptation bandwidth, *σ^max^* was constant or differed by spectral context. Estimating a separate *σ* for each range condition significantly contributed to overall explained variance for N1 (Likelihood ratio test comparing models with constant or varying *σ*: *χ*^2^(2)=16.4, p=0.0003), but not for P2 (*χ*^2^(2)=1.9, p=0.38). Nevertheless, we decided to fit a model with separate *σ^max^* to both the N1 and P2 data for comparison purposes. We found that, interestingly, the values of *σ^max^* were consistently close to range/4 (Figure 5C, diagonal dashed line) which is the mean interval between neighboring tones on the sequence-specific frequency axis (since there are 5 possible tone frequencies and 4 frequency intervals between them).

To additionally test whether the modulation of *σ* by range was significant, we performed 101 bootstrap repetitions (sampling with replacement from the 31 participants) and analyzed the resulting *σ^max^* using an LME model (fixed factors: Intercept and range, random factors: Intercept and range grouped by bootstrap number, Methods). Table 3 shows that the dependence of the bandwidth on the range was highly significant with large effect sizes, for both N1 and P2. Thus, although introducing separate bandwidths for the different ranges did not improve the overall fit of the model to the P2 data, nevertheless the bandwidths estimated separately for each range were highly stable. In particular, this test provides evidence that even for the P2 data, the adaptation bandwidth at the smallest range was significantly smaller than at medium and large range (Figure 5C, Table 3).

**Table 3.** LME results – effect of frequency range on adaptation bandwidth. The data consisted of 101 bootstrap estimates of adaptation bandwidth (sampling with replacement from the 31 participants of Experiment 3 and repeating parameter estimation procedure) for each range condition and potential type (606 data points). The model consisted of fixed and random factors (grouped by bootstrap number) for each of the listed predictors (Methods). Column headers are similar to Tables 1 and 2.

| Predictors |  | Estimate | SE | F(1,602) | p-value | d |
| --- | --- | --- | --- | --- | --- | --- |
| Intercept | N1 | 2.7 | 0.37 | 52 | 1.7E-12 | 0.72 |
|  | P2 | 5.7 | 0.39 | 218 | 2.7E-42 | 1.5 |
| Range | N1 | 0.22 | 0.013 | 283 | 2E-52 | 1.7 |
|  | P2 | 0.13 | 0.013 | 95 | 5.8E-21 | 0.97 |

The dependence of *σ^max^* on the stimulus frequency range could potentially be caused merely by an overall reduction of peak amplitudes in sequences with smaller ranges, without any relation to the specific order of the tones in the sequence. To test if this is the case, we permuted the order of data trials within each range condition separately, and repeated the parameter estimation procedure 100 times. The values of *σ^max^* obtained using the permuted data showed a slight modulation by frequency range (Figure 5C, pale dashed lines) but it was not significant (slope of *σ^max^* as a function of frequency range estimated for permuted data, N1: 0.04, 95% CI=(−0.03 0.11), not different from 0: t(298)=0.99, p=0.32, P2: 0.05, 95% CI=(−0.02, 0.12), t(298)=1.37, p=0.17). Furthermore, the slopes of *σ^max^* as a function of frequency range were significantly larger when using the real and bootstrapped data compared to using the permuted data (Figure 5C) both for N1 (slopes as a function of range: 0.21, 95% CI=(0.19 0.24), note that this confidence region did not overlap with the confidence region of the slope for permuted data presented above) and for P2 (0.13, 95% CI=(0.1 0.15)). Thus, the overall reduction of amplitudes in sequences with smaller ranges was not sufficient to explain the spectral context effect.

## DISCUSSION

Our results suggest neural sensitivity to past stimulation at two distinct time scales. Specifically, we show that a longer integration time scale (~3 s) is evident in the ERPs early after stimulus onset, at a latency of about 100 ms (N1), and is reflected in apparent sensitivity to the mean tone frequency. A shorter integration time scale (~1 s) is evident at a longer latency after stimulus presentation, about 200 ms (P2) and is reflected in sensitivity to the immediately preceding stimulus. Further, we show quantitatively that both of these response components can be accounted for by frequency-specific adaptation of neural populations having distinct time constants of recovery. Finally, by varying the frequency range of the stimulus sequence we show that spectral context affects the frequency adaptation bandwidth of these neural populations, independent of integration times.

### Context integration across several time scales

The N1 and P2 auditory evoked potentials were among the first recorded human EEG responses (Davis, 1939), but their neural generators and the computations underlying these responses are still not fully understood (Picton, 2011; Lanting et al., 2013), although their generators are thought to reside mainly in auditory cortex. N1 is well-known for being strongly attenuated by stimulus repetition, an effect termed adaptation, habituation or refractoriness (Crowley and Colrain, 2004; Picton, 2011). Previous studies have established that the time scale of N1 adaptation is longer than about 1 s (Zacharias et al., 2012; Okamoto and Kakigi, 2014; Herrmann et al., 2016). N1 adaptation in a sequence of pure tones was well accounted for by the frequency-specific adaptation model we used here (Herrmann et al., 2013, 2014).

In contradistinction, the effect of stimulus repetition on P2 is more controversial. P2 was sometimes suggested to be less affected by adaptation than N1 (Crowley and Colrain, 2004; Herrmann et al., 2013, 2016) and sometimes more (Lanting et al., 2013). Importantly, previous studies failed to account for P2 responses using adaptation models (Herrmann et al., 2013, 2016), including exactly the same frequency-specific adaptation model we used here (Herrmann et al., 2013). However, Herrmann et al. (2013) used a single, predetermined, time constant for both N1 and P2 when fitting the model to the data (1.8 s, based on Sams et al., 1993). Here, by developing a methodology to directly estimate the time constant from the data, we show that both N1 and P2 responses can be well explained by frequency-specific adaptation mechanisms, but that P2 has a shorter time scale of adaptation than N1. Our results contribute to characterizing the functional distinction between the N1 and P2 responses and thus support the claim that they are generated by distinct neural populations (Knight et al., 1980; Hari et al., 1982; Lanting et al., 2013).

The time constants we report may depend on specifics of the paradigm we used. For example, the sequences we used did not contain long-term information over time scales longer than about 5 seconds. For this reason, we only tested recovery time constants up to 5 seconds. In Experiment 1 the time constant estimated for N1 reached the maximal value of 5 seconds (Figure 5A). This may indicate that the time scale of N1 sensitivity may be even longer than what we report here. Indeed, other human electrophysiology studies used sequences that contained longer-term structure and reported longer time scales of integration, ~10 s, for N1 (Herrmann et al., 2016). Furthermore, Costa-Faidella et al. (2011) found that in pure tone sequences, P2 and the MMN were sensitive both to regularities established across short (< 1 second) and long (~10 seconds) time scales. It could be that sequences embedding longer-term structure can reveal longer time scale sensitivity of P2. However, the long-term structure in Costa-Faidella et al. was the violation of a globally established regularity. Thus, the reported long-term sensitivity of P2 may reflect deviance-detection and prediction mechanisms, whereas the effects we report here may be due to adaptation mechanisms that are not specifically related to regularity extraction.

### Early processing of long and later processing of short time scales

Interestingly, we found that the earlier response component, N1, had a longer time constant compared to P2, which peaks on the scalp later. Similarly, early auditory processing of long-scale sequence properties and later processing of short-scale properties was recently reported in a MEG study (Maheu et al., 2019). These results are surprising because longer time scales of integration are frequently associated with a higher-level of processing, which is expected to take place later in the processing hierarchy. For example, studies employing ‘global-local’ paradigms found that early sensory cortex was sensitive to short-term, local regularities whereas later and more widespread activity was sensitive to long-term regularities (Bekinschtein et al., 2009; Dürschmid et al., 2016). However, the distinction between short- and long-term regularities in the latter studies, established in the context of deviance detection paradigms, are likely to reflect neural mechanisms that are distinct from the ones underlying our results, obtained in a passive paradigm with no regularities or deviance.

One can interpret our results in terms of resolution, associating a longer scale with coarser temporal resolution and a shorter scale with a higher resolution that allows to process finer details. This view corresponds with theories such as the frame-and-fill model developed for vision (Bar, 2006; Snyder et al., 2012), or reverse hierarchy theory initially also developed for vision (Ahissar and Hochstein, 2004) but later extended to audition (Nahum et al., 2008). By these theories, we quickly obtain a general gist of the sensory scene at a coarse resolution, requiring long-scale averaging. Later on, the perceptual system may gradually resolve the details, expressed in the shorter temporal scales of processing at later stages, consistent with our results.

### Sensitivity to the mean

Temporal averaging plays an important role in perception (Hollingworth, 1910; Haberman et al., 2009; Albrecht and Scholl, 2010; McDermott and Simoncelli, 2011; McDermott et al., 2013). Our results manifest modulation of neural responses due to the mean of previous stimuli: N1 responses were most adapted for stimuli closest to the mean, consistent with previous reports (Ulanovsky et al., 2004; Herrmann et al., 2013, 2014). Although we did not measure perception in this study, the neural machinery described here may be related to the neural computation of mean and its influence on perception. For instance, judgements in a frequency discrimination task were biased towards the mean of past stimuli (Raviv et al., 2012; Lieder et al., 2019). This perceptual bias to the mean was correlated to ERP adaptation (Jaffe-Dax et al., 2017), potentially linking it with the neural effects we study here. We suggest that the mean is represented by the adaptation level of a frequency-specific, yet wideband, neuronal population that adapts momentarily to stimuli as they come, then recover with a long time constant. The frequency corresponding to the most adapted population is the mean frequency of the sequence. This results in a time-dependent estimate of the mean frequency, which effectively averages past stimuli in a sliding window whose duration depends on the time constant of recovery.

### Frequency bandwidth rescales to spectral range

We found that adaptation bandwidths rescale to the range of frequency distributions in the sequence. Our results are consistent with previous reports for N1 (Herrmann et al., 2013, 2014, 2015) and generalize them for P2 as well. This suggests that sensitivity of adaptation bandwidths to the spectral context is a general feature of auditory cortex, independent of the integration time scales. Somewhat similar effects, although in the context of deviance detection, have been shown by Garrido et al. (2013).

What could be the mechanism of this sensitivity to the spectral context? Dynamic adjustment of neural input-output functions due to changes in statistical properties of preceding stimulation has been directly shown using neural measurements in non-human animals for various perceptual dimensions such as light intensity (Dunn and Rieke, 2006), visual motion (Brenner et al., 2000) and whisker motion in the rat barrel cortex (Maravall et al., 2007). In the auditory modality, rescaling of neuronal tuning to varying statistical properties of sounds were reported in the bird midbrain (Nagel and Doupe, 2006), mammalian inferior colliculus (Kvale and Schreiner, 2004; Dean et al., 2008; Dahmen et al., 2010) and primary auditory cortex (Blake and Merzenich, 2002; Gourévitch et al., 2009; Rabinowitz et al., 2011). Thus, the change of neural response patterns due to stimulation statistics could reflect activity from the same neural population that undergoes rapid changes to network connectivity (Arnsten et al., 2010; Rabinowitz et al., 2011). Alternatively, the changes of neuronal response characteristics we measured following different stimulation ranges may also reflect recruitment of distinct neuronal populations, each with its own fixed, distinct spectral and temporal response properties (Lee et al., 2016; Osman et al., 2018).

In conclusion, we find distinct responses in the auditory cortex which are sensitive to the context over shorter and longer time scales, and which quickly and automatically adjust to the spectral context. We show that the temporal effects are likely due to different rates of release from adaptation. The mechanism of the spectral effect remains to be determined.

## Abbreviations

ERP: Event-related potentials
SOA: stimulus-onset asynchrony
*LL^max^*: maximum log-likelihood
ms: milliseconds
s: seconds

## Acknowledgements

We thank research assistants who aided with EEG data acquisition and preprocessing: Michal Rabinovits, Eden Krispin, Anael Benistri. TIR was supported by the Hoffman Leadership and Responsibility Program at the Hebrew University. LYD was supported by the Israel Science Foundation and Jack H. Skirball research fund. IN was supported by a grant from the Israel Academy of Sciences (390/13) and by AdERC grant 340063 (project RATLAND), and is the Milton and Brindell Gottlieb Chair in Brain Science. The funders had no role in study design, data collection and analysis, decision to publish, or preparation of the manuscript.

## Author contributions

TIR formulated the study, conducted experiments, developed and conducted analysis methods, and wrote the first draft of the paper. GM conducted experiments. IN and LYD jointly supervised the study, and participated in the design of the experiments, formulation and application of analysis methods and writing the paper.

## Declaration of interests

The authors declare no competing interests. LYD declares he is a co-founder and advisor of InnerEye Ltd.

## Data and Code Availability

Data and code are available via Open Science Framework (OSF): http://osf.io/mswhv

## SUPPLEMENTARY FIGURES AND TABLES

**Supplementary Figure S1.**
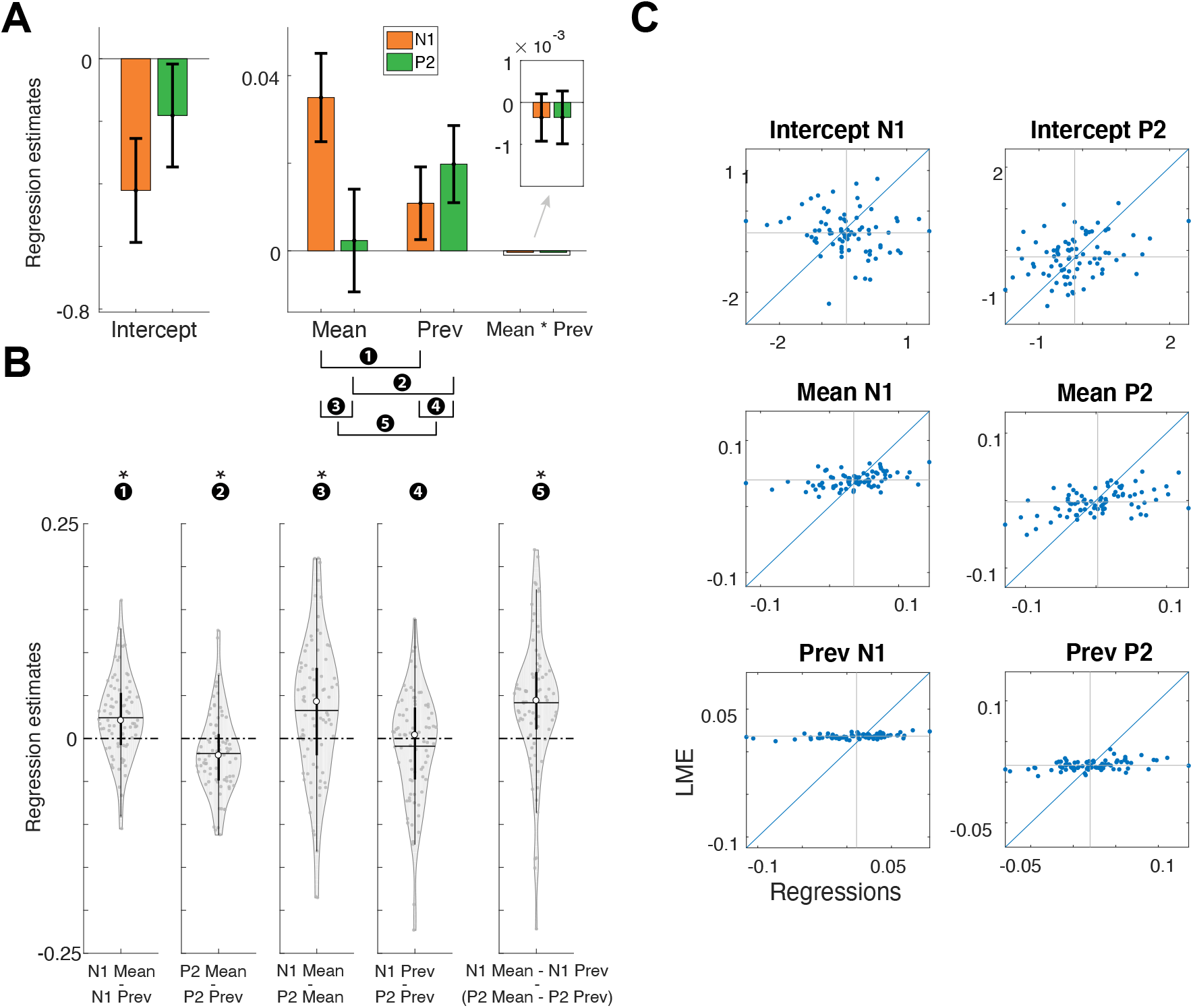
Second-level analysis of the estimates from single-participant regressions. Compare with the results of the linear mixed effects model (LME) in Figure 2. **A** and **B** are the same type of plots as in Figure 2, but using the estimates from single-participant regressions instead of the LME estimates. For the regressions, the same predictors were used as in the fixed factors of the LME. The error bars in A represent the 95% confidence intervals calculated across the group of participant-specific estimates. Each dot in **B** represents the difference between the relevant estimates per each participant. **Horizontal lines represent the mean, white circles the medians and thick and thin black vertical lines represent the 25% and 75% percentiles, respectively**. **C** – Scatter plots of LME vs. regression estimates per participant. Participant-specific estimates in the LME were calculated by adding the common fixed-effect estimate to the participant-specific random estimate. The fact that LME estimates are closer to the mean value than regression estimates illustrates the tendency of the LME to constrain the variability across participants.

**Supplementary Table S1.** Second-level analysis of participant-specific regression estimates, for the long- and short-term context predictors – ‘Interval-Mean’ and ‘Interval-Previous’. N1 and P2 amplitudes (voltage in μV was standardized using a z-score transform) modeled using the 8 predictors as in Table 1. Here a simple linear regression was run separately for each participant (only fixed, no random factors). For each predictor the estimates for all 79 participants were collected. First row, columns 2 to 5: Estimate – mean estimate across participants. SE – standard error across the group. Cohen’s d – mean estimate/SD. P-value – calculated from a t-test comparing the estimate to 0. Five bottom lines refer to a paired t-test comparing between groups of estimates. X vs. Y stands for a subtraction between X and Y. In parenthesis - values from the corresponding LME model (Table 1) for comparison, when relevant.

| Predictors |  | Estimate | SE | t(78) | p-value | d |
| --- | --- | --- | --- | --- | --- | --- |
| Intercept | N1 | -0.42<br>(-0.51) | 0.084<br>(0.078) | -5 | 2.9E-6<br>(8.8E-11) | -0.57<br>(-0.74) |
|  | P2 | -0.18<br>(-0.81) | 0.083<br>(0.074) | -2.2 | 0.03<br>(0.016) | -0.24<br>(-0.27) |
| Interval-Mean | N1 | 0.035<br>(0.04) | 0.005<br>(0.004) | 7 | 1.1E-9<br>(6.7E-17) | 0.78<br>(0.95) |
|  | P2 | 0.002<br>(-0.002) | 0.006<br>(0.005) | 0.4 | 0.69<br>(0.69) | 0.04<br>(-0.04) |
| Interval-Previous | N1 | 0.01<br>(0.02) | 0.004<br>(0.004) | 2.6 | 0.01<br>(2.2E-7) | 0.29<br>(-0.59) |
|  | P2 | 0.02<br>(0.02) | 0.004<br>(0.004) | 4.4 | 2.5E-5<br>(1.7E-8) | 0.5<br>(0.64) |
| Interval-Mean *<br>Interval-Previous | N1 | -0.0004<br>(-0.0008) | 0.0003<br>(0.0002) | -1.2 | 0.2<br>(0.001) | -0.14<br>(-0.36) |
|  | P2 | -0.0003<br>(-0.0002) | 0.0003<br>(0.0002) | -1.1 | 0.26<br>(0.33) | -0.12<br>(-0.11) |

| Comparisons between pairs of predictors | t(78) | p-value | d |
| --- | --- | --- | --- |
| N1 vs. P2 Interval-Mean | 3.5 | 7.1E-4<br>(2.6E-9) | 0.39<br>(0.67) |
| N1 vs. P2 Interval-Previous | -1.2 | 0.21<br>(0.6) | -0.12<br>(-0.05) |
| N1 Interval-Mean vs. -previous | 4.5 | 2.2E-5<br>(4.7E-7) | 0.5<br>(0.57) |
| P2 Interval-Mean vs. -previous | -3.3 | 1.3E-3<br>(3.7E-6) | -0.37<br>(-0.52) |
| N1 Interval-Mean vs. -previous<br>vs.<br>P2 Interval-Mean vs. -previous | 4.7 | 9.4E-6<br>(1E-11) | 0.53<br>(0.77) |

**Supplementary Figure S2.**
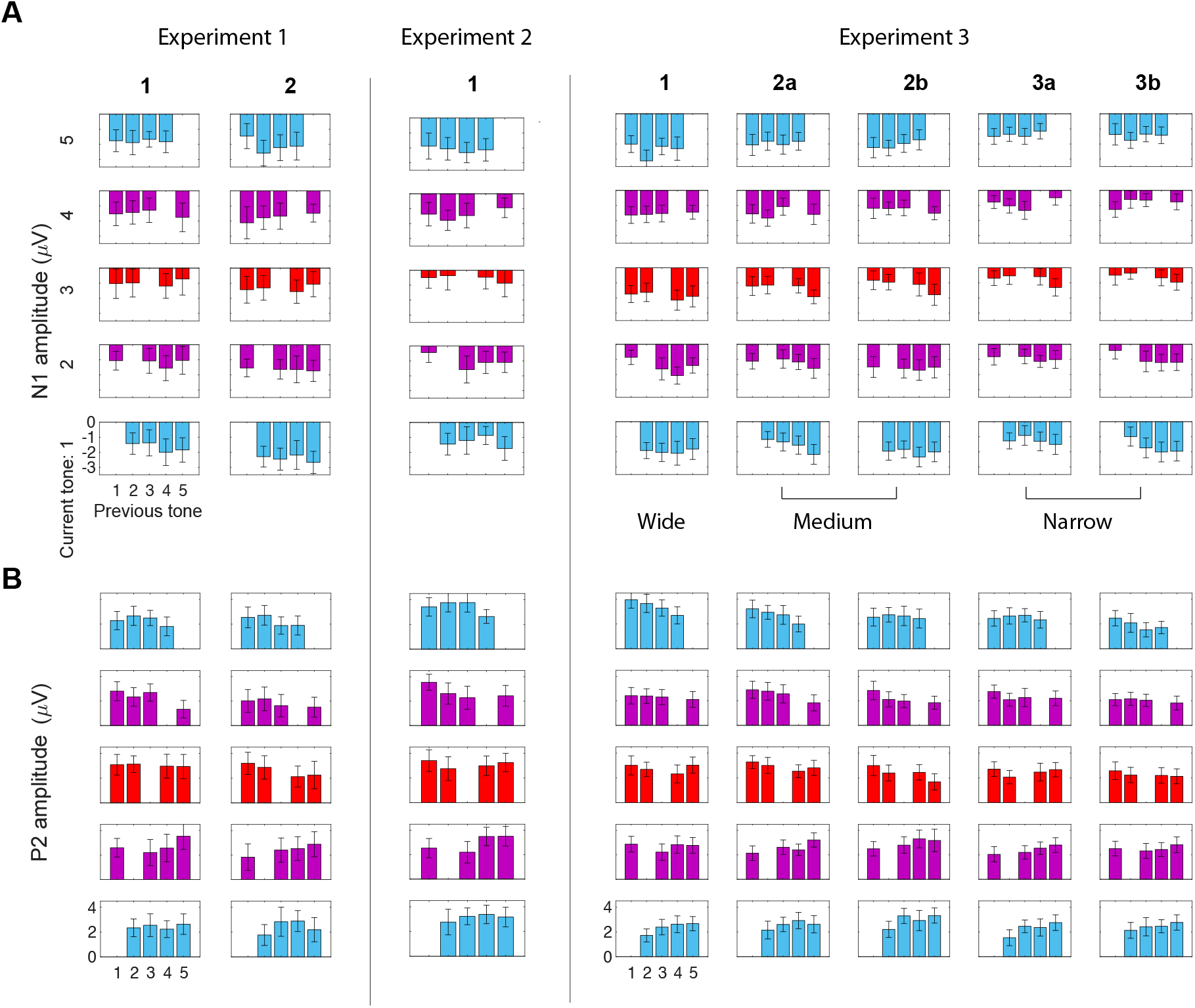
Average N1 and P2 amplitudes, all combinations of current and previous tones. Bar graphs depict average N1 (**A**) or P2 (**B**) peak amplitudes averaged across participants for every pair of current and previous tones, in every experiment and block type. Error bars represent 95% confidence intervals of the peaks across participants. Numbers under experiment titles denote block types. Tones are ranked from 1 to 5 from low to high frequency, within each block type (as illustrated in Figure 1A). Each row of bar graphs shows the N1/P2 amplitudes for a specific tone rank contingent on the previous tone rank. Since there were no immediate repetitions of the same tone, each graph misses the bar for the immediate repetition (e.g. tone 5 following tone 5 of a sequence, tone 4 following tone 4, etc.). For specific tones’ frequencies see Figure 1B and methods.

**Supplementary Table S2.**
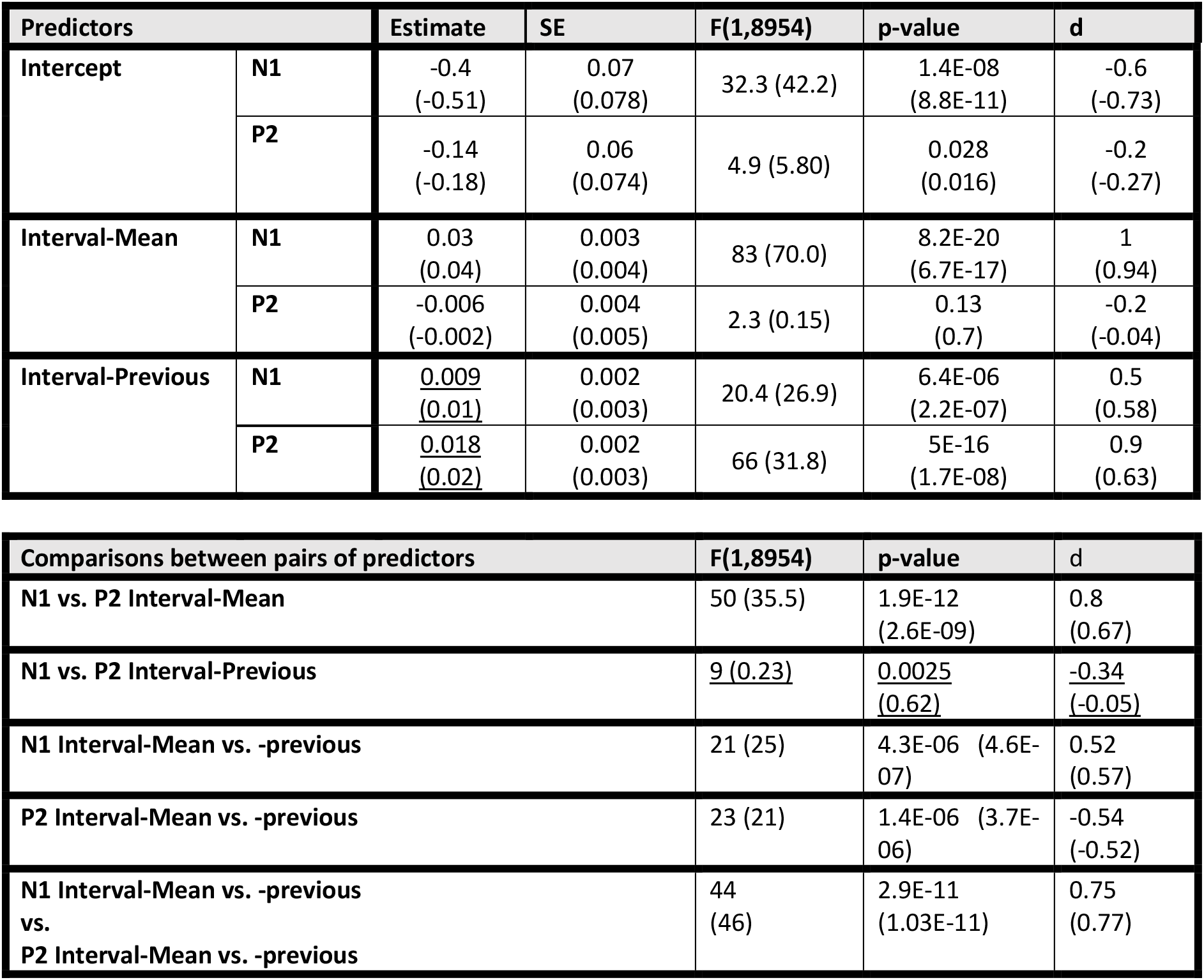
Linear mixed effects (LME) results for the long- and short-term context predictors – ‘Interval-Mean’ and ‘Interval-Previous’, excluding their interaction term. Entries are similar to Table 1. The actual values of Table 1 (from the LME that included the interaction term) are added in parenthesis for comparison. Note that the main difference between these results and those presented in Table 2 is that in this case the contribution of the short-term context variable (Interval-Previous) to N1 (but not P2) was reduced, and as a result the contrast between Interval-Previous contributions to N1 and P2 is larger and significant (highlighted with underline).

| Predictors |  | Estimate | SE | F(1,8954) | p-value | d |
| --- | --- | --- | --- | --- | --- | --- |
| Intercept | N1 | -0.4<br>(-0.51) | 0.07<br>(0.078) | 32.3 (42.2) | 1.4E-08<br>(8.8E-11) | -0.6<br>(-0.73) |
|  | P2 | -0.14<br>(-0.18) | 0.06<br>(0.074) | 4.9 (5.80) | 0.028<br>(0.016) | -0.2<br>(-0.27) |
| Interval-Mean | N1 | 0.03<br>(0.04) | 0.003<br>(0.004) | 83 (70.0) | 8.2E-20<br>(6.7E-17) | 1<br>(0.94) |
|  | P2 | -0.006<br>(-0.002) | 0.004<br>(0.005) | 2.3 (0.15) | 0.13<br>(0.7) | -0.2<br>(-0.04) |
| Interval-Previous | N1 | <u>0.009</u><br><u>(0.01)</u> | 0.002<br>(0.003) | 20.4 (26.9) | 6.4E-06<br>(2.2E-07) | 0.5<br>(0.58) |
|  | P2 | <u>0.018</u><br><u>(0.02)</u> | 0.002<br>(0.003) | 66 (31.8) | 5E-16<br>(1.7E-08) | 0.9<br>(0.63) |

| Comparisons between pairs of predictors |  | F(1,8954) | p-value | d |
| --- | --- | --- | --- | --- |
| N1 vs. P2 Interval-Mean |  | 50 (35.5) | 1.9E-12<br>(2.6E-09) | 0.8<br>(0.67) |
| N1 vs. P2 Interval-Previous |  | <u>9 (0.23)</u> | <u>0.0025</u><br><u>(0.62)</u> | <u>-0.34</u><br><u>(-0.05)</u> |
| N1 Interval-Mean vs. -previous |  | 21 (25) | 4.3E-06 (4.6E-07) | 0.52<br>(0.57) |
| P2 Interval-Mean vs. -previous |  | 23 (21) | 1.4E-06 (3.7E-06) | -0.54<br>(-0.52) |
| N1 Interval-Mean vs. -previous<br>vs.<br>P2 Interval-Mean vs. -previous |  | 44<br>(46) | 2.9E-11<br>(1.03E-11) | 0.75<br>(0.77) |

**Supplementary Table S3.** Linear mixed effects (LME) results including interactions with frequency range and the interaction term Interval-Mean*interval_previous. This Table is almost identical to Table 2, but the LME model included the interaction terms Interval-Mean * Interval-Previous for N1 and P2. Note that the Interval-Mean * Interval-Previous terms had no significant contribution (highlighted with underline) and therefore were omitted from the LME model reported in Table 2. However, they affected the statistical comparisons between estimates (lower Table). In parenthesis – values from Table 2 (when relevant) for comparison.

| Predictors |  | Estimate | SE | F(1,6186) | p-value | d |
| --- | --- | --- | --- | --- | --- | --- |
| Intercept | N1 | -0.76<br>(-0.76) | 0.11<br>(0.1) | 52<br>(55) | 6.3E-13<br>(1.2E-13) | -1.3<br>(-1.4) |
|  | P2 | -0.39<br>(-0.36) | 0.12<br>(0.11) | 10.7<br>(9.7) | 0.001<br>(0.002) | -0.60<br>(-0.57) |
| Interval-Mean | N1 | 0.054<br>(0.054) | 0.01<br>(0.01) | 17.2<br>(17.3) | 3.3E-5<br>(3.2E-5) | 0.76<br>(0.76) |
|  | P2 | -0.029<br>(-0.028) | 0.014<br>(0.014) | 4.5<br>(4.2) | 0.035<br>(0.041) | -0.39<br>(-0.37) |
| Interval-Previous | N1 | 0.027<br>(0.027) | 0.009<br>(0.009) | 8.7<br>(8.8) | 3.2E-3<br>(3.0E-3) | 0.54<br>(0.54) |
|  | P2 | 0.042<br>(0.043) | 0.009<br>(0.009) | 19.7<br>(21.2) | 9.0E-6<br>(4.1E-6) | 0.81<br>(0.84) |
| Interval-Mean * Interval-Previous | N1 | <u>0.00002</u> | <u>0.0005</u> | <u>0.0014</u> | <u>0.97</u> | <u>0.01</u> |
|  | P2 | <u>0.0005</u> | <u>0.0005</u> | <u>1.3</u> | <u>0.25</u> | <u>0.21</u> |
| Range * Intercept | N1 | 0.019<br>(0.018) | 0.004<br>(0.003) | 17.5<br>(36.2) | 2.9E-5<br>(1.9E-9) | 0.76<br>(1.1) |
|  | P2 | 0.008<br>(0.0041) | 0.004<br>(0.003) | 3.07<br>(1.8) | 0.08<br>(0.18) | 0.32<br>(0.24) |
| Range * Interval-Mean | N1 | -0.0011<br>(-0.0011) | 0.0004<br>(0.0004) | 6.08<br>(7.2) | 0.014<br>(0.007) | -0.45<br>(-0.49) |
|  | P2 | 0.0004<br>(0.0006) | 0.0004<br>(0.0004) | 0.97<br>(2.5) | 0.32<br>(0.11) | 0.18<br>(0.29) |
| Range * Interval-Previous | N1 | -0.0006<br>(-0.0006) | 0.0003<br>(0.0003) | 3.2<br>(4) | 0.073<br>(0.047) | -0.33<br>(-0.36) |
|  | P2 | -0.0009<br>(-0.0007) | 0.0003<br>(0.0003) | 7.5<br>(6.2) | 6.0E-3<br>(1.2E-2) | -0.50<br>(-0.46) |

| Comparisons between pairs of predictors | F(1,6186) | p-value | d |
| --- | --- | --- | --- |
| N1 vs. P2 (Range * Intercept) | 2.95<br>(10.9) | 0.086<br>(0.0009) | 0.31<br>(0.6) |
| N1 vs. P2 (Range * Interval-Mean) | 5.96<br>(9) | 0.015<br>(0.002) | -0.45<br>(-0.55) |
| N1 vs. P2 (Range * Interval-Previous) | 0.46<br>(0.13) | 0.5<br>(0.72) | 0.12<br>(-0.07) |

**Supplementary Figure S3.**
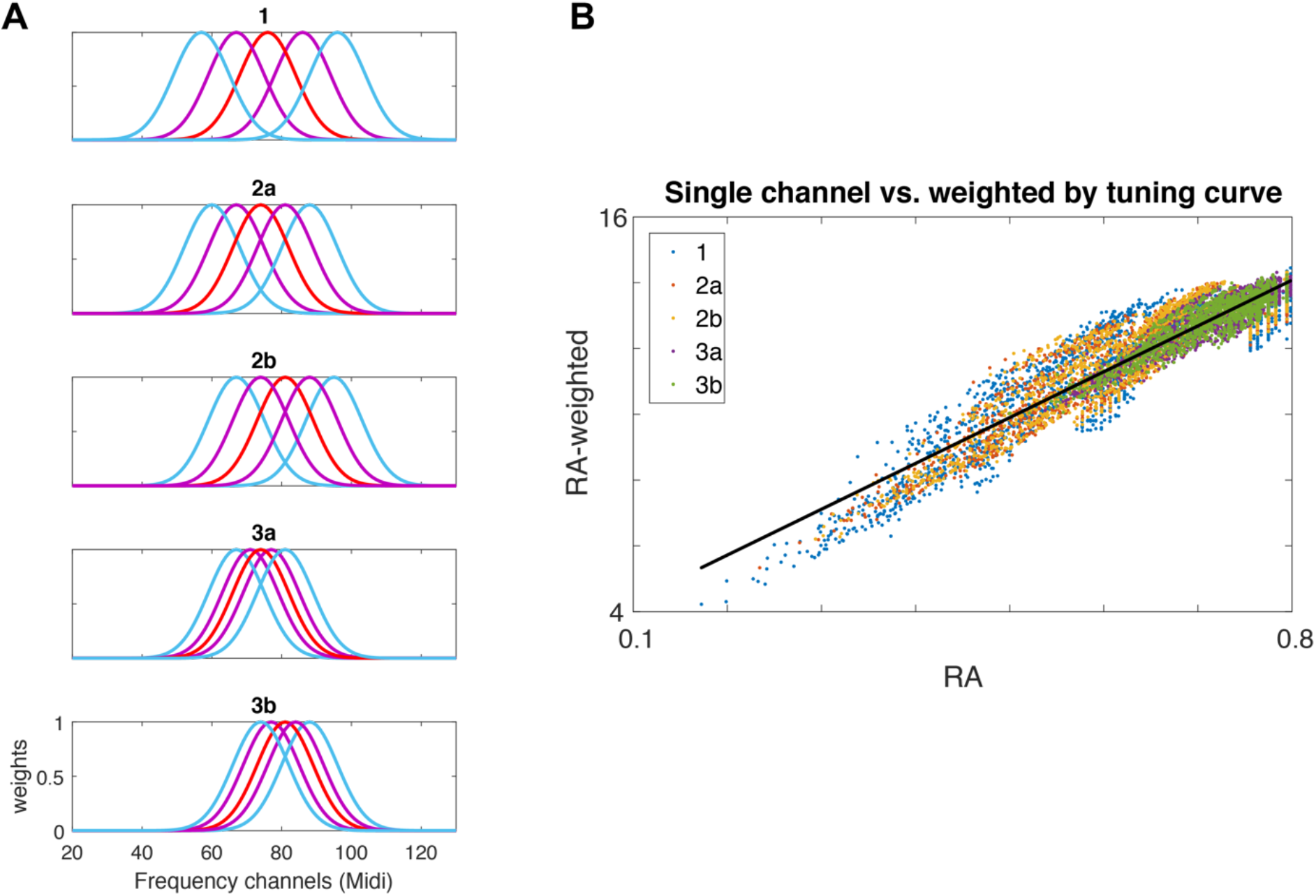
Comparison of RA (the response adaptation model used in this work) to RA weighted by tuning curves. See Methods for a definition of RA. **A** – Gaussian tuning curves. Each plot corresponds to a specific block type in Experiment 3 (specified in the titles). In each plot, each Gaussian describes the relevant weights for summing up contributions from neural populations with best frequencies displayed along the x-axis. The 5 Gaussians correspond to the computed responses to each of the 5 tones, color code similar to Figure 1. **B** - Scatter plot of the weighted RA values (y axis) versus single channel RA values (x axis) as used in this work following Herrmann et al. (2013, 2014, 2015).

**Supplementary Figure S4.**
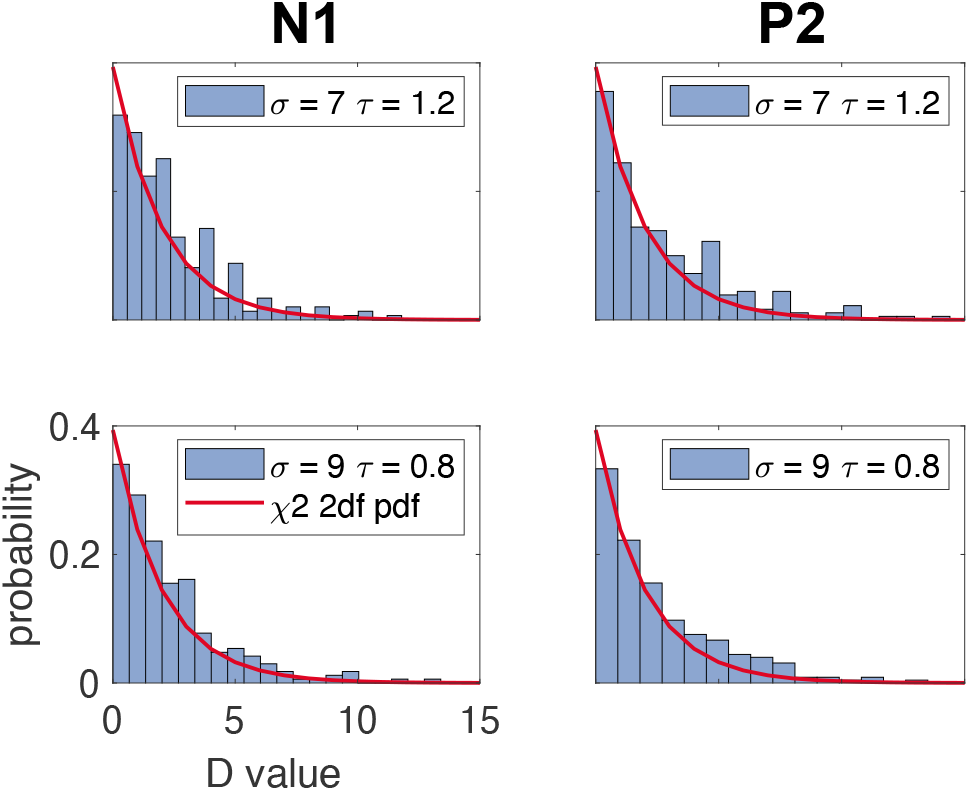
Comparison of the null D distribution to chi-square with 2 degrees of freedom. Blue - histograms of null D distribution computed for 2 representative values of 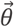: σ = 7 semitones and τ = 1.2 seconds (upper) or σ = 9 semitones and τ = 0.8 seconds (lower) using N1 (left) or P2 (right) data. Null distribution was simulated by fitting the model to scrambled data, such that the order of data trials did not match that of model predictions, 250 times. The red curve is the theoretical χ^2^ distribution with 2 degrees-of-freedom.

**Supplementary Figure S5.**
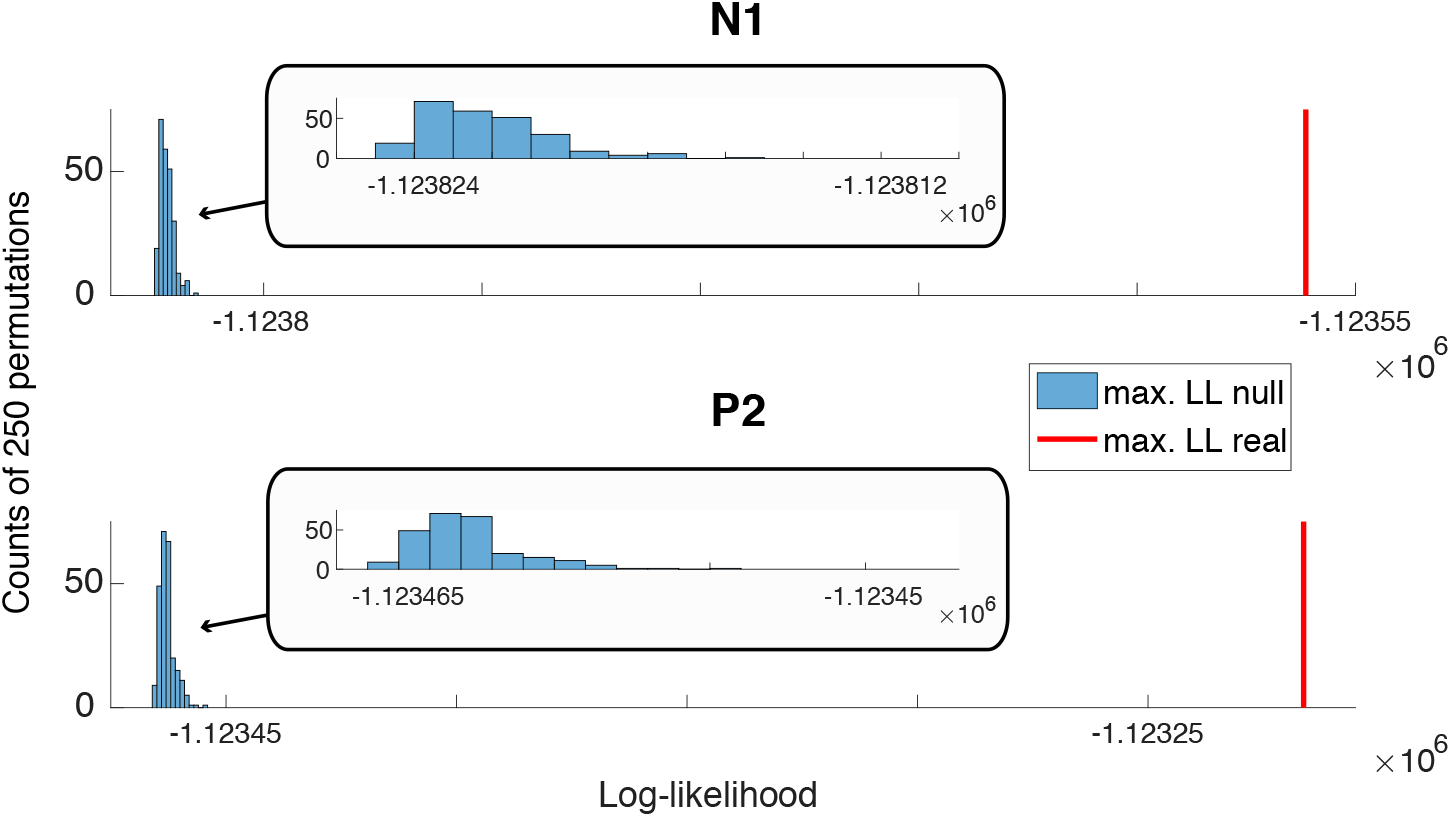
General significance of model fit. **Blue bars** - Histograms of null distributions of the maximal Log-Likelihood (LL) of LME model fit. Histograms are constructed using 250 repetitions of model fit after permuting the order of data trials, using the data from all experiments. **Red lines** – the maximal value of LL out of all tested 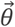 values (the LL value at 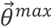), when fitting the model using the real (non-scrambled) data. **Top** – The model fitted to N1 data trials. **Bottom** – The model fitted to P2 data trials. **Insets** – Magnifying x-axis values.

**Supplementary Figure S6.**
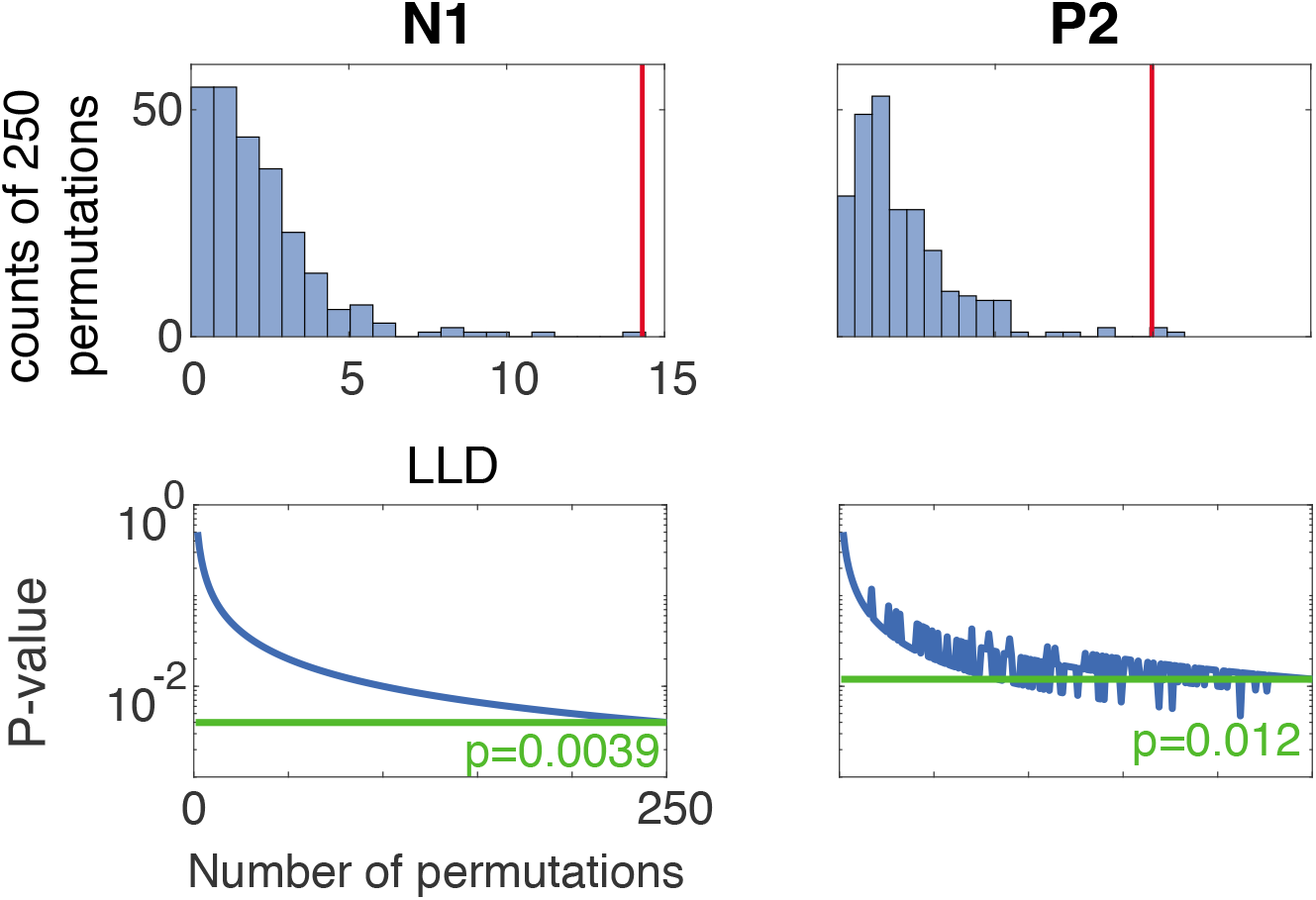
Differences between log-likelihood at τ^max^ and τ^max^ of other potential type (LLD) relative to null distribution. Null distribution was calculated by fitting the model and estimating parameters for 250 repetitions of random scrambling of data trials such that their order did not match model predictions. **Top** – blue histograms illustrate the null LLD distribution calculated using the 250 repetitions and red lines indicate the real LLD. **Bottom** – The green displayed value of p is calculated using all 250 repetitions for the null distribution. The blue line illustrates the p-value as a function of the number of repetitions used (scramble groups are selected randomly from the pool of 250 scramble repetitions calculated). The green line denotes the p-values calculated when using 250 repetitions. Note that the p-values seem to be an upper bound of the real p-value. **Left** - N1. **Right** - P2.

## Notes

http://osf.io/mswhv

### Competing Interest Statement

The authors have declared no competing interest.

### Summary of Updates

New revision following a peer-review

